# A new genome-scale metabolic model of oleaginous microalgae with refined lipid metabolism elucidates *Microchloropsis gaditana* mutant phenotypes

**DOI:** 10.1101/2023.12.06.570374

**Authors:** Clémence Dupont-Thibert, Sylvaine Roy, Sónia Carneiro, Bruno Pereira, Rafael Carreira, Paulo Vilaça, Séverine Collin, Eric Maréchal, Elodie Billey, Gilles Curien, Maxime Durot, Juliette Jouhet

**Affiliations:** Laboratoire de Physiologie Cellulaire et Végétale, CNRS, CEA, INRAE, Univ. Grenoble Alpes, 17 Avenue des Martyrs, 38000 Grenoble, France; TotalEnergies, OneTech, Centre de Recherche de Solaize CRES, Chemin du Canal, 69360 Solaize, France; SilicoLife Lda. Rua do Canastreiro, 15 4715-387 Braga, Portugal

**Author notes:** Corresponding author Juliette Jouhet.

**Keywords:** genome-scale metabolic modeling, lipid metabolism, microalgae, *Microchloropsis gaditana*

## Abstract

The oleaginous microalga *Microchloropsis gaditana* (formerly *Nannochloropsis gaditana*) has gained large interest due to its potential to produce lipids for a wide range of biotechnological applications. To optimize *M. gaditana* growth conditions and develop new strains to enhance lipid synthesis and accumulation, a broad understanding of the organism metabolism is essential. Computational models such as genome-scale metabolic models constitute powerful tools for unravelling microorganism metabolism. In this work we present iMgadit23, a new genome-scale metabolic model for *M. gaditana*. Model covers 2330 reactions involving 1977 metabolites and associated with 889 genes. Pathways involved in membrane and storage glycerolipid biosynthesis and degradation have undergone thorough manual curation and have been comprehensively described based on current knowledge of *M. gaditana* lipid metabolism. Additionally, we developed a detailed 2D-pathway map of model content to provide a systems-level visualization of *M. gaditana* metabolism. We demonstrated the predictive capabilities of iMgadit23, validating its ability to qualitatively and quantitatively capture in vivo growth phenotypes under diverse environmental and genetic conditions. Model was also able to capture the role of the Bubblegum acyl-CoA synthetase in remodeling *M. gaditana* lipid metabolism. iMgadit23 and its 2D map constitute valuable tools to increase understanding of *M. gaditana* metabolism and deciphering mutant phenotypes, specifically in the context of lipid metabolism. The model holds significant promise in predicting *M. gaditana* metabolic capabilities, facilitating strain engineering, and optimizing cultivation processes for a broad range of industrial applications.

## Introduction

Microalgae are photosynthetic microorganisms that have attracted significant attention from the biotechnology industry because of their ability to efficiently transform renewable resources (CO_2_ and light) into biomass and other valuable molecules such as lipids. Due to the wide range of lipids applications in diverse fields such as biofuels, green chemistry, health or food (Chu, 2012; Rizwan et al., 2018; Spolaore et al., 2006), ongoing research involves the metabolic engineering of various microalgae strains and the optimization of culture processes to enhance lipid synthesis and accumulation (Fu et al., 2019; Muñoz et al., 2021; Rock et al., 2021; Sajjadi et al., 2018). Oleaginous microalgae were selected for this purpose such as diatoms or the Nannochloropsis clade, now divided into *Nannochloropsis* and *Microchloropsis* genera (Fawley et al., 2015). However, by difference with green algae, the metabolism of these microalgae is not well known due to their complex evolutionary history deriving from multiple endosymbioses (Keeling, 2004; Petroutsos et al., 2014). Due to this complex evolutionary history, a wide variety of lipid compositions and lipid metabolism pathways have been reported within microalgal diversity (Kong et al., 2018; Li-Beisson et al., 2019). The first visible impact of *Nannochloropsis* and *Microchloropsis* evolution path is the presence of a single secondary plastid with four limiting membranes resulting from a primary and a secondary endosymbiosis (Fawley et al., 2015; Petroutsos et al., 2014). In nitrogen starvation, these microalgae accumulate triacylglycerols (TAGs) up to 40 % on a dry weight basis (Simionato et al., 2013). They also produce eicosapentaenoic acid (EPA, 20:5 20 carbon atoms: five unsaturations), a very long-chain polyunsaturated fatty acid (VLC-PUFA) which has been shown to provide many health benefits to humans (Al-Hoqani et al., 2017; Ma et al., 2016).

The lipid molecules identified in both *Nannochloropsis* and *Microchloropsis* genera are similar (Mühlroth et al., 2017). Numerous gaps of knowledge regarding lipid metabolism pathways and their regulation need to be filled for these genera and more generally for microalga strains containing plastids with four limited membranes (Hoffmann and Shachar-Hill, 2023; Li-Beisson et al., 2019). Filling these gaps is crucial to enhance lipid synthesis and accumulation. Inclusion of accurate and species-specific description of lipid metabolic pathways into a genome-scale metabolic model (GEM) would constitute a valuable tool to investigate lipid metabolism further and to guide the production of these biotechnological targets. Derived from genome annotations and experimental data, GEMs are metabolic networks that collect all known metabolic information of a biological system, including the metabolites, reactions, genes, enzymes and associated gene-protein-reaction (GPR) rules. Constraint-Based Reconstruction and Analysis (COBRA) methods perform systems-level analyses on GEMs to simulate how genetic and environmental factors affect phenotype, represented by flux distributions (Heirendt et al., 2019). For example, Flux Balance Analysis (FBA) is widely used to predict the phenotype under certain culture conditions by calculating an optimal network state that optimizes an objective function (Orth et al., 2010). Biomass Objective Functions (BOFs), consuming all metabolites required to produce one gram of dry weight biomass (gDW), are commonly used as FBA objectives to simulate microorganisms growth. The resulting solution from FBA is typically not unique, i.e., multiple flux distributions can lead to the same growth rate. A common approach to decrease the possible solution space of FBA is to use the parsimonious version of FBA (pFBA). This approach finds a flux distribution with minimum absolute flux values among the alternative optima, assuming that the cell attempts to achieve its objective while allocating the minimum amount of resources (Lewis et al., 2010). In the last 25 years, COBRA methods have proven to be powerful tools for understanding and redesigning the metabolism of microbial strains including microalgae (Gu et al., 2019; Tibocha-Bonilla et al., 2018; Ye et al., 2022).

Given their potential for industrial biotechnological applications, efforts have been done in the last few years to reconstruct GEMs of *Nannochloropsis* and *Microchloropsis* species. The first draft metabolic model of *Nannochloropsis* sp. was published by (Pham, 2016). It was followed in 2017 by the publication of two fully reconstructed models of *M. gaditana* (iRJ1321) and *M. salina* (iNS937) (Loira et al., 2017; Shah et al., 2017). Finally, a model of *M. gaditana* automatically reconstructed using AuCoMe (Automated Comparison of Metabolism) was recently published (Belcour et al., 2023). Among these four GEMs, only iRJ1321 and iNS937 allow biomass production. In the Pham model, biomass reaction was added based on Viridiplantae data. However, not all precursors for this BOF can be synthesized by the model, making growth simulation impossible (Pham, 2016). The Belcour model, originally reconstructed for conducting phylogenetic comparisons between species rather than predicting flux distributions, lacks any BOF. Even if iRJ1321 and iNS937 can be used to predict flux distributions, they are sub-optimal choices for the study of lipid metabolism due to the presence of non-functional or mis-compartmentalized pathways, and to missing components. In this study, we present iMgadit23, a new metabolic model for *M. gaditana* with a detailed description and modeling of lipid metabolism. This model was used to simulate growth behavior of the strain in different media and genetic conditions, and to unravel the metabolic phenotype of the acsbg mutant strain, which has an interesting lipid profile, notably with an eight-fold increase in TAG accumulation compared to the wild-type strain. (Billey et al., 2021). By validating the simulation to the experimental data, we can propose iMgadit23 to the researcher community as a useful tool to visualize the impact of a mutation on metabolism, to test the putative lethality of a genetic knock-out or even to assess which metabolic pathways to modify in order to reach a targeted biomass composition.

## Results

iMgadit23: a new genome-scale metabolic model for the oleaginous microalga **M. gaditana**.

### General model properties

GEM model was automatically reconstructed based on the genome sequence of *M. gaditana* B-31, before being manually curated and refined. The final curated model, iMgadit23, consists of 889 genes, 2330 reactions and 1977 metabolites (1155 chemical entities, i.e. when excluding metabolites duplicated in multiple compartments). The model includes eight compartments: extracellular (denoted by “e” in the model), cytosol (c), chloroplast stroma (h) and lumen (l), endoplasmic reticulum (ER, r in model), peroxisome (p), mitochondrial matrix (m) and intermembrane space (IMS, i in model) (Figure1A). Model compartments were defined to encompass the intra-organelle space plus its membrane enclosure. For example, ER compartment covers both ER membrane and lumen. Most metabolites (36%) and reactions (30%) are located in the cytosol. While the chloroplast stroma contains 21% of all metabolites and reactions, the ER contains 24% of all reactions and 14% of all metabolites. All of the 2330 reactions are classified into 15 different subsystems mainly based on Kyoto Encyclopedia of Genes and Genomes (KEGG) pathway database (Kanehisa and Goto, 2000). These subsystems cover central and specialized metabolism (Figure1B). Subsystems were divided into more detailed sub-subsystems available in iMgadit23 notes.

Given the promising lipid accumulation capacity of *M. gaditana* and its biotechnological potential, membrane and storage glycerolipid biosynthesis and degradation pathways were described in detail in iMgadit23. In total, lipid metabolism encompasses 1017 reactions (44% of all model reactions). Among them, 600 reactions, localized either in plastid, cytosol, or ER are involved in lipid biosynthesis (fatty acid biosynthesis, glycerolipid metabolism and steroid biosynthesis). The remaining 417 reactions are involved in fatty acids (FAs) degradation (β-oxidation in mitochondria and peroxisome) and glycerolipids degradation (TAG, MGDG and PC) in cytosol, plastid and ER (Figure1C).

Central metabolism as modeled in iMgadit23 is represented in Figure 2. An exhaustive 2D-pathway map was drawn and is available as JSON file compatible with Escher (https://github.com/Total-RD/Mgaditana-GEM) (King et al., 2015). This map provides a systems-level representation of *M. gaditana* metabolism, and constitutes a valuable tool to visualize and analyze biological data including reaction data (metabolic fluxes), metabolite data (metabolomics), and genomic data (transcriptomics data).

**Figure 1.**
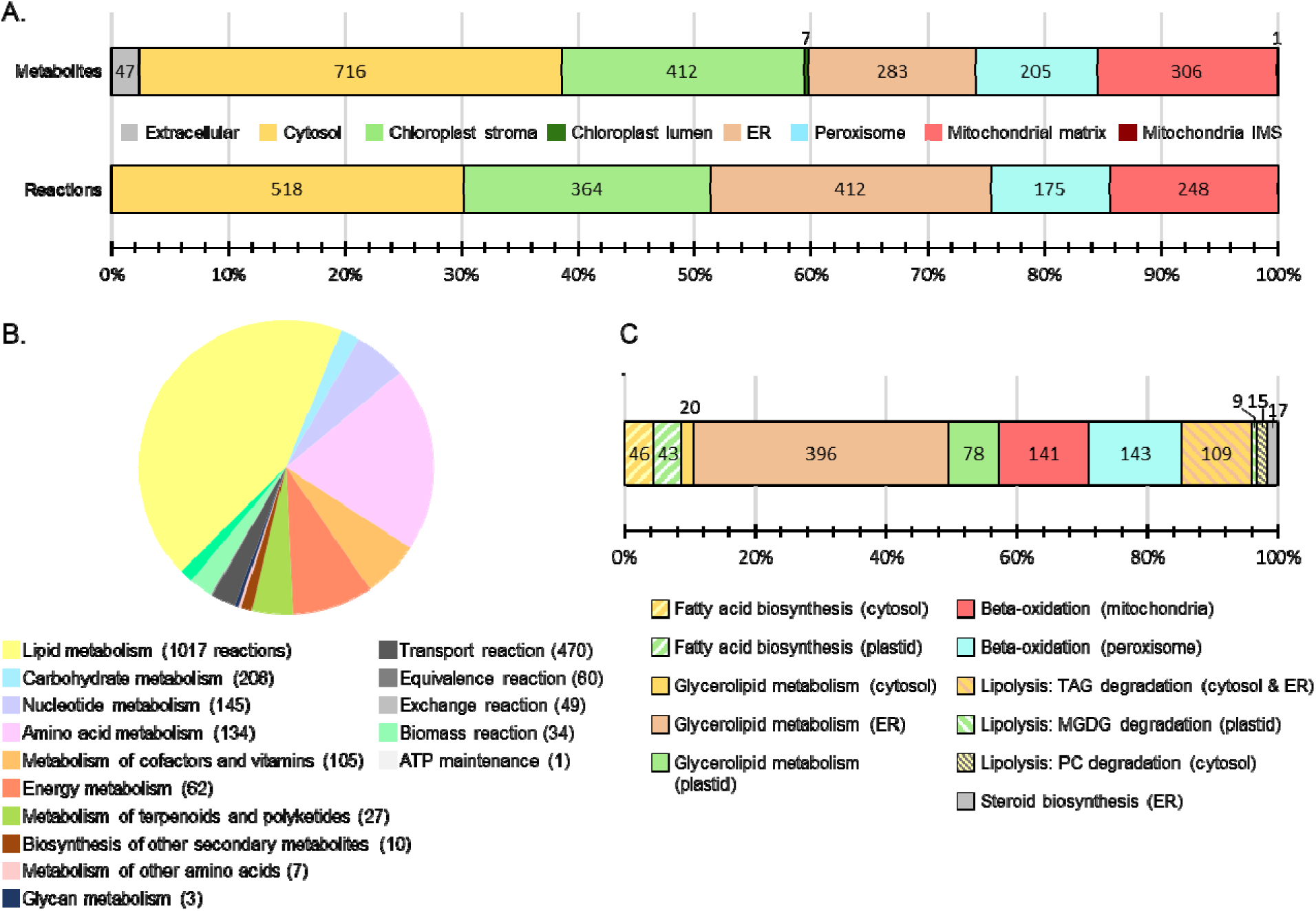
General properties of iMgadit23 metabolic model. (A) Distribution of metabolites and metabolic reactions in the eight compartments: extracellular, cytosol, chloroplast stroma and lumen, endoplasmic reticulum (ER), peroxisome, mitochondrial matrix and intermembrane space (IMS). Transport, Exchange, Equivalence and Biomass reactions included in the model are not represented. (B) Distribution of reactions into 15 subsystems (KEGG metabolic pathways), number of reactions in each subsystem is indicated between parentheses. (C) Distribution of lipid metabolism reactions by compartmentalized pathways; compartments are indicated between parentheses.

**Figure 2.**
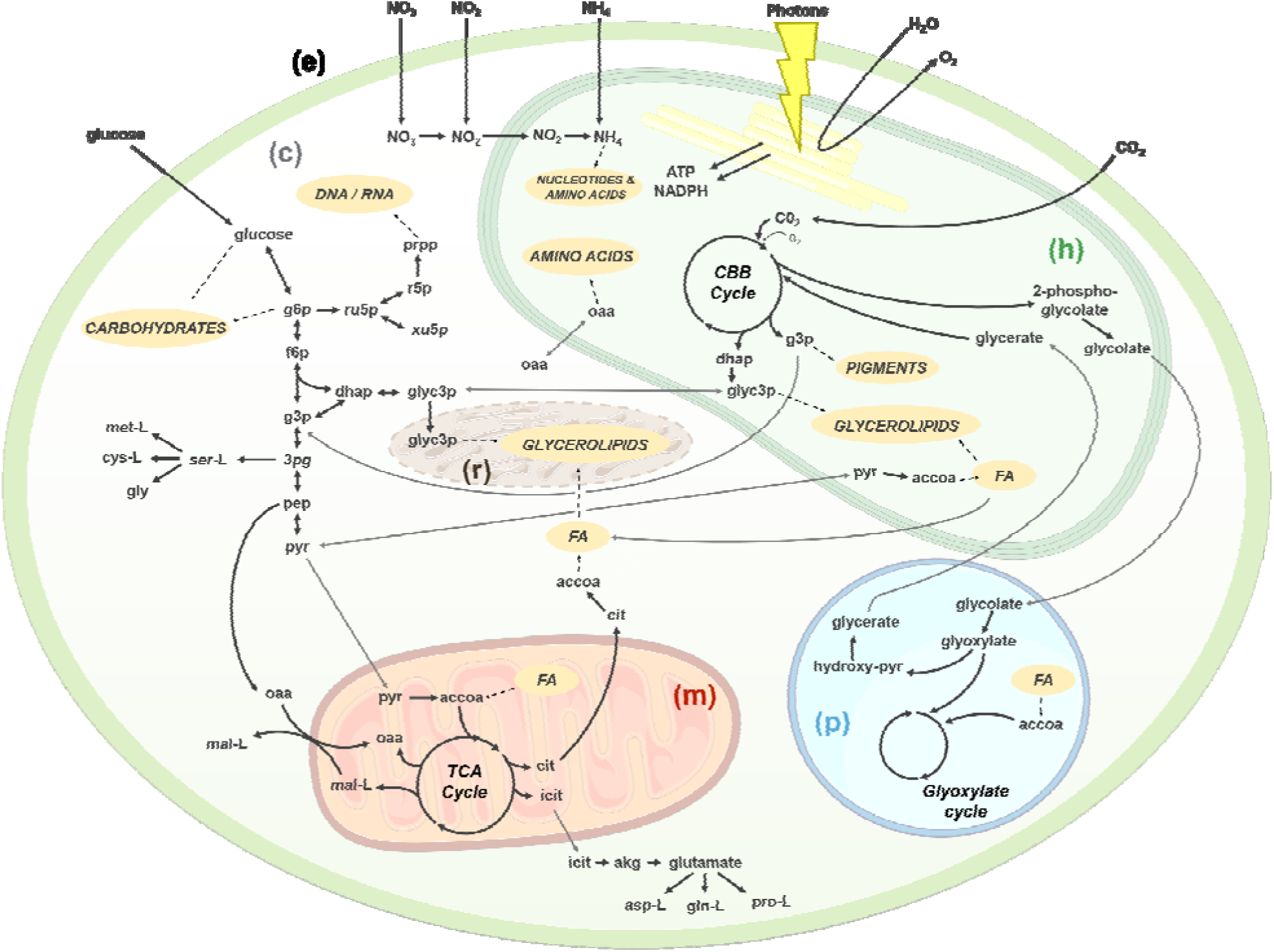
Central metabolism as modeled in iMgadit23. The main model compartments are represented: cytosol (c), chloroplast stroma (h), extracellular medium (e), mitochondria matrix (m), peroxisome (p) and ER (r). Metabolites: 3-phosphoglycerate (3pg), acetyl-CoA (accoa), L-aspartate (asp-L), α ketoglutarate (akg), citrate (cit), L-cysteine (cys-L), fatty acids (FA), fructose-6-phosphate (f6p), glucose-6-phosphate (g6p), glyceraldehyde-3-phosphate (g3p), L-glutamine (gln-L) glycerol-3-phosphate (glyc3p), glycine (gly), dihydroxyacetone phosphate (dhap), isocitrate (icit), malate (mal-L), methionine (met-L), oxaloacetate (oaa), phosphoenolpyruvate (pep), phospho-ribose 1-diphosphate (prpp), proline (pro-L), pyruvate (pyr), ribose 5-phosphate (r5p), ribulose 5-phosphate (ru5p), serine (ser-L), xylulose 5-phosphate (xu5p). Pathways: Calvin–Benson–Bassham (CBB) cycle, tricarboxylic acid cycle (TCA) cycle.

### Photosynthesis

Photosynthesis occurs in two steps of coupled reactions. (1) The “light phase” produces reducing power in the form of NADPH and chemical energy in the form of ATP. (2) In the “dark phase”, both ATP and NADPH are subsequently used in the Calvin–Benson–Bassham (CBB) cycle to produce glyceraldehyde-3-phosphate (G3P) from inorganic carbon (CO_2_).

#### Linear Electron Flow

In the “light phase”, light energy (photons) is used to extract electrons from water, generating O_2_ as a by-product. Electrons are transported through thylakoid-bound proteins including the photosystem II (PSII), the cytochrome b_6_/f complex (cyt b_6_/f) also known as plastoquinol-plastocyanin reductase, and the photosystem I (PSI). Electrons are then used to reduce ferredoxin and finally NADP^+^. This electron transfer process is denoted as linear electron flow (LEF). In iMgadit23, LEF is modeled by four reactions: Photosystem II (PSII_h), Plastoquinol/Plastocyanin Reductase (PPR_h), Photosystem I (PSI_h), Ferredoxin oxidoreductase (FDXO_h). LEF is coupled to the movement of protons (H^+^) into the internal space of the thylakoids (lumen) creating a proton motive force, the driving force for ATP synthesis by ATP synthase (ATPSYN_h) (Supplemental Results, Figure S1A).

#### Photosynthesis yields

Examination of the CBB cycle shows that assimilation of one CO_2_ molecule requires consumption of three ATP and two NADPH. In LEF, electron transfer results in six H^+^ translocated to the lumen every two electrons transferred from water to NADP^+^. According to this stoichiometry, production of two NADPH by LEF requires four electrons, resulting in 12 H^+^ translocated to the lumen (Curien et al., 2016). Based on these calculations, in our model we assumed a coupling factor of H^+^/ATP = 12/3 for chloroplastic ATP synthase, meaning that 12 H^+^ are required to produce three ATP. This assumption results in an ATP/NADPH production ratio of 3/2, meeting CBB demand. Real value of the ATP/NADPH ratio produced by LEF is still debated, but is probably lower than 1 (Allen, 2002). Moreover coupling factor of plastidial ATP synthase was reported to have a H^+^/ATP ratio of 14/3, resulting in a lower production of ATP (ATP/NADPH = 1.286) (Hahn et al., 2018).

Cyclic electron flow (CEF) has been proposed as a complementary pathway to increase ATP production by the photosynthetic chain (Allen, 2002). Thus, in the “light phase” of photosynthesis, LEF would produce two NADPH leading to 12H^+^ translocation to chloroplast lumen, extra 2H^+^ required to produce 3 ATP would be translocated through CEF requiring two extra photons. CEF is not described in iMgadit23. To overcome this limitation, we introduced the distinction between absorbed photons (photon_abs) and photons used for LEF (photon_lef) in iMgadit23, concomitantly with the addition of a pseudo-reaction: PSQUANTUM_h, that artificially converts 10 absorbed photons to produce 8 photons LEF. The addition of this reaction allows easy modulation of absorbed photon/ATP ratio.

pFBA simulations with BOF as objective function were performed with different absorbed photon/ATP ratios, highlighting the significant impact of this ratio on model predictions (Supplemental Results, Figure S1B) Indeed, absorbed photons being the only limiting “substrate” in our simulations, when increasing the absorbed photon/ATP ratio from 8/3 to 10/3 the maximum growth rate predicted by iMgadit23 is decreased by 20%. Similar impacts on ATP and NADPH production were also predicted by iMgadit23 (Supplemental Results, Figure S1C and D).

pFBA approach, using BOF maximization as objective function was performed to estimate quantum yield (QY), defined as number of moles of CO_2_ fixed per mole of photon absorbed. For this simulation, maximum photon uptake rate was constrained to 19 mmol .gDW^-1^.h^-1^. iMgadit23 predicts a QY of 0.047 mol .mol ^-1^, which is consistent with the maximum experimental QY of 0.05 mol .mol ^-1^ measured for *Nannochloropsis* sp. (Raso et al. 2012). Without PSQUANTUM pseudo-reaction, iMgadit23 predicts a QY of 0.063 mol .mol ^-1^. These results illustrate that addition of PSQUANTUM allows to better represent photosynthesis efficiency in iMgadit23. Moreover, absorbed photons/LEF photons ratio being easily changed, this pseudo-reaction constitutes a first step to investigate photosynthesis efficiency in *M. gaditana*.

### Lipid metabolism

First, a standardized nomenclature of metabolite and reaction identifiers was introduced in iMgadit23. Lipid nomenclature is detailed in Supplemental Material and Methods, Table S3. Then, based on current knowledge of lipid metabolism of *M. gaditana* and other photosynthetic organisms, FAs and glycerolipids biosynthesis and degradation were added to iMgadit23. These pathways are schematized in Figure 3 and represent lipid metabolism diversity in terms of glycerolipid class and acyl chains profiles (Jouhet et al., 2017). To the author’s knowledge, iMgadit23 represents the most up-to-date and accurate GEM of *Nannochloropsis* and *Microchloropsis* both in term of reaction content and localization.

**Figure 3.**
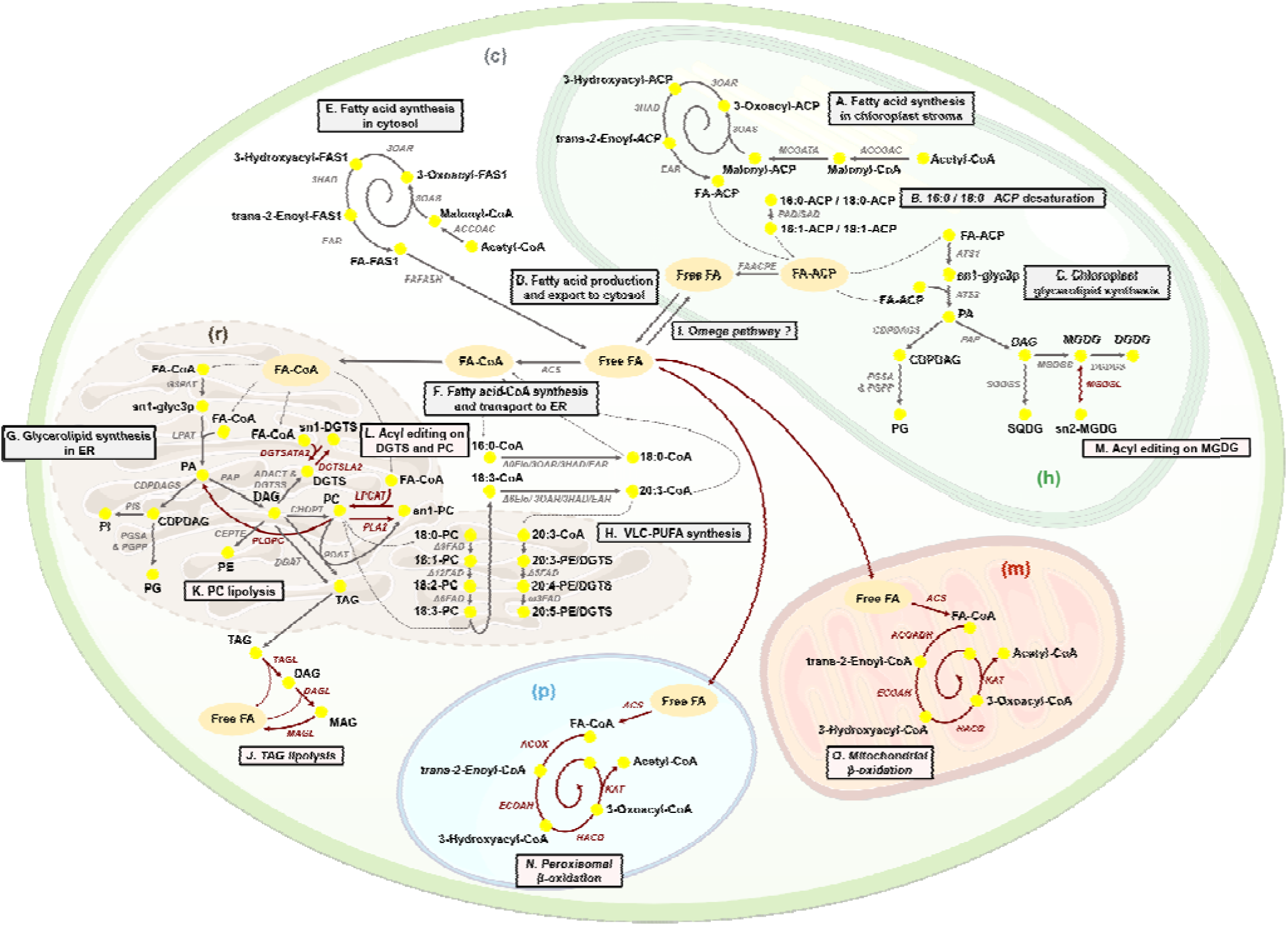
Lipids metabolism as modeled in iMgadit23. Grey and red arrows represent respectively lipid synthesis (A to I); lipid degradation and acyl editing (L to O). (A) Fatty acids (FAs) are initially synthetized in chloroplast stroma, starting with an ATP-dependent carboxylation of acetyl-CoA by an acetyl-CoA carboxylase (ACCOAC), producing malonyl-CoA which is subsequently converted into malonyl-acyl carrier protein (ACP) by the malonyl CoA-ACP malonyltransferase (MCOATA). Then, 3-oxoacyl-ACP synthase III catalyzes the condensation of malonyl-ACP with acetyl-ACP (3OAS). 3-oxoacyl-ACP is reduced by a 3-oxo-acyl reductase (3OAR) into 3-hydroxyacyl-ACP which serves as a substrate for an enoyl-dehydratase. 2,3-trans-enoyl-ACP produced is finally reduced by an enoyl-reductase (EAR) into an acyl-ACP containing two additional carbons compared to the initial substrate, up to a chain length of 14, 16 or 18 carbons. (B) Saturated FAs 16:0-ACP and 18:0-ACP can be desaturated by a stromal acyl-ACP desaturase forming 16:1-ACP and 18:1-ACP (PAD161_h and SAD181_h reactions). The FA-ACPs have two fates: they can (C) be acylated to glycerol-3-phosphate (glyc3p) by glycerol-3-phosphate acyl-ACP acyltransferases (ATS1 and ATS2) to produce precursors of four glycerolipids species: PG, SQDG, MGDG and DGDG (respectively via CDPDAG synthase (CDPDAGS), PG phosphate synthase (PGSA), PG phosphate phosphatase (PGPP), phosphatidic acid phosphatase (PAP), SQDG Synthase (SQDGS), MGDG Synthase (MGDGS) and DGDG Synthase (DGDGS)), or (D) be converted to free FAs by an acyl-ACP hydrolase (FAACPE) and exported to the cytosol. (E) FAs can also be de novo synthesized and elongated in cytosol. (F) Cytosolic free FAs can be thio-esterified to CoA by acyl-CoA synthetase (ACS) and (G) be used in ER as initial precursors for synthesis of TAG via the Kennedy pathway by the glycerol 3-phosphate acyltransferase (G3PAT), the lysophosphatidic acid acyltransferase (LPAT), the PAP, the phospholipid:diacylglycerol acyltransferase (PDAT) and the diacylglycerol acyltransferase (DGAT). Betaine lipid DGTS is formed by a 2-step reaction of DAG with S-adenosylmethionine (ADACT and DGTSS). The membrane phosphoglycerolipids PI, PG, PE, and PC are respectively synthetized via CDPDAGS, PI Synthase (PIS), PGSA, PGPP, ethanolamine phosphotransferase (CEPTE) and choline phosphotransferase (CHOPT). (H) Δ0-ELO, and Δ6-ELO elongases as well as Δ9, Δ12, Δ6, Δ5 and Δ15/ω3 fatty acyl desaturase (FAD) are required for very-long-chain polyunsaturated fatty acids (VLC-PUFAs) synthesis. (I) VLC-PUFAs 20:4 and 20:5 are then imported into the plastid via the “omega pathway”. (J) TAGs can be degraded via TAG-, DAG-, and MAG-lipases. (K) PC lipolysis can occur via Phospholipase D (PLDPC). (L) FAs can be edited on PC by Phospholipase A2 (PLA2) and lyso-PC acyl transferase (LPCAT) and DGTS can be edited by DGTS lipase A2 (DGTSLA2) and DGTS acyl transferase A2 (DGTSATA2). (M) Acyl-editing on MGDG (MGDG lipase (MDGDL)). Free FAs can be degraded via β-oxidation both in peroxisome (N) and mitochondria (O). Model compartments are denoted (c) cytosol, (h) chloroplast stroma, (m) mitochondrial matrix, (p) peroxisome and (r) ER.

#### Lipid biosynthesis

FAs bound to acyl carrier proteins (ACPs) are de novo synthetized by the multi-subunit bacterial type II FA synthase (FASII) complex in chloroplast stroma (Figure 3A), generating essentially 16:0-ACP and 18:0-ACP which can undergo desaturation by a stromal acyl-ACP desaturase, leading to the production of 16:1-ACP and/or 18:1-ACP (Figure 3B). FA-ACPs can either be used to esterify glycerol-3-phosphate (glyc3p in iMgadit23) to produce precursors for four glycerolipids: phosphatidylglycerol (PG), sulfoquinovosyldiacylglycerol (SQDG), monogalactosyldiacylglycerol (MGDG) and digalactosyldiacylglycerol (DGDG) (Figure 3C), or be converted into free FAs to be exported to the cytosol (Figure 3D) (Li-Beisson et al., 2019). FA elongation pathway by FASI complex was also added in iMgadit23 cytosol (Figure 3E). Indeed, transcription of the cytosolic type I FAS was reported in *M. gaditana* in stress conditions such as high light exposition, suggesting possibility of cytosolic de novo FA synthesis (Alboresi et al., 2016). Cytosolic free FA are then thioesterified to CoA by acyl-CoA synthetases (ACSs), forming acyl-CoAs which are then used as precursors for all membrane phosphoglycerolipids in the ER i.e., phosphatidylinositol (PI), phosphatidylglycerol (PG), phosphatidylethanolamine (PE), phosphatidylcholine (PC); triacylglycerol (TAG) and the betaine lipids diacylglyceryl-3-O-40-(N,N,N-trimethyl)-homoserine (DGTS) (Figure 3F and G) (Li-Beisson et al., 2019). 16:0-CoA is also the initial substrate for a succession of elongations and desaturations forming at first 18:0-CoA and ending up with very-long-chain polyunsaturated FAs (VLC-PUFAs) up to 20:5 (EPA) (Figure 3H) (Dolch et al., 2017; Vieler et al., 2012). Re-importation of VLC-PUFAs into the plastid is required for the assembly of MGDG, DGDG, SQDG and PG. This process, known as the “omega pathway”, remains to be fully characterized (Figure 3I) (Dolch et al., 2017; Petroutsos et al., 2014).

#### Lipolysis and acyl editing

Since growth and survival under fluctuating environmental conditions require permanent remodeling or turnover of membrane lipids as well as rapid mobilization of storage lipids, lipid degradation and acyl editing are important processes in microalgae. The process of acyl editing refers to a de-acylation of an acyl group from a given lipid and re-acylation on another lipid. It allows for fatty acid desaturation and modification prior to removal and further incorporation into other glycerolipids. Most of what is currently known about glycerolipid degradation and acyl editing has been established in plants and much less is known in algae (Hoffmann and Shachar-Hill, 2023; Kong et al., 2018). Few data are available for *M. gaditana* and other *Nannochloropsis* and *Microchloropsis* species. Based on current knowledge for *M. gaditana*, algae and plants, TAG, DAG and MAG (monoacylglycerol) lipolysis (Figure 3J) and the hydrolysis of PC by Phospholipase D, producing PA and a free choline group (Figure 3K) were added to our model. Acyl editing on PC, DGTS and MGDG substrates were also added (Figure 3L and M). Reactions addition regarding glycerolipid degradation and acyl editing are further justified in Supplemental Material and Methods.

#### β-oxidation

In *Nannochloropsis* and *Microchloropsis* species, FA degradation via β-oxidation may occur both in peroxisome and mitochondrion (Figure 3N and O) (Li-Beisson et al., 2019; Poliner et al., 2015). It begins with the dehydrogenation of acyl-CoA to trans-2-enoyl-CoA either via the reduction of dioxygen (O_2_) to generate peroxide (ACOX in peroxisome) or via the reduction of FAD to FADH2 (ACOADH in mitochondria). In both compartments, trans-2-enoyl-CoA is then hydrated, dehydrogenated, and cleaved into an acetyl-CoA and an acyl-CoA, two-carbons shorter compared to the initial substrate. For unsaturated FAs, two additional enzymes are required, cis-Δ3-Enoyl CoA isomerase that converts a cis-Δ3 bond to a trans-Δ2 bond (32ECOAI reactions) and the 2,4 Dienoyl CoA reductase that reduces 2,4-dienoyl-CoA into trans-Δ3-enoyl-CoA (24E2COAR reactions).

#### Lipid metabolites and reactions identifiers

Due to the limitations of BiGG identifiers for lipid metabolites (Witting, 2020), a new nomenclature has been established in iMgadit23 to facilitate the understanding of lipid metabolism. Lipids are identified by their class acronym and detailed acyl chain composition following sn-1, sn-2 (and sn-3 for TAGs) order. Acyl chains are labeled according to the format “number of carbons:number of double bonds in acyl chain”. For examples, hexadecanoic acid also known as palmitic acid or FA 16:0 is identified as “c160”, and TAG 16:0/18:1/16:1 is identified as tag160_181_161. Detailed lipid nomenclature is available in Supplemental Material and Methods, Table S3.

Due to the symmetry of the TAG chemical structure, the TAG molecule sn-1 sn-2 sn-3 is identical to the TAG molecule sn-3 sn-2 sn-1, which is not the case in iMgadit23 where tag_sn1_sn2_sn3 and tag_sn3_sn2_sn1 constitute distinct metabolites due to their distinct synthesis pathways. To overcome this limitation, “equivalence” pseudo-reactions were added to the model, converting identical TAGs into a unique pseudo-TAG. For example, to represent equivalence between tag160_181_161 and tag161_181_160, two reactions converting respectively these TAGs into pseudo-TAG tag502 (50 carbons and 2 unsaturations) were added to the model (reactions were named EQformula_TAG502_160181161_c and EQformula_TAG502_161181160_c). In total iMgadit23 contains 24 “EQformula_TAG” reactions. Same modeling methodology was applied for all lipid classes.

### Biomass Objective Function

In GEMs, a BOF consumes resources essential for the growth of a given microorganism, weighted to represent mole fraction of each component in one gram of dry weight. In iMgadit23 BOF contains stoichiometric coefficients for 121 metabolites, including 78 lipids. These metabolites are consumed by six pseudo-reactions representing the different macromolecules (PROTEINS, DNA, RNA, LIPIDS, CARBOHYDRATES and PIGMENTS). Ten pseudo-reactions representing glycerolipid classes (TAG, DAG, PC, PE, PI, PG, DGTS, DQDG, MGDG and DGDG) were also added. Detailed lipidomic data, in terms of glycerolipid profiles and FA profiles in glycerolipid classes, of *M. gaditana* WT grown under photoautotrophy were used to calculate stoichiometric coefficients of lipids (Billey et al., 2021). In fine, biomass composition in iMgadit23: CH_1.8_O_0.49_N_0.11_P_0.005_S_0.003_Mg_0.001_, is consistent with the molecular formula of microalgae cell reported in literature: CH_1.83_O_0.48_N_0.11_P_0.01_ (Ma et al., 2022). For detailed composition of BOF see Supplemental Material and Methods and Supplemental Files S1 and S2.

In addition to biomass composition, ATP requirements for both growth-associated maintenance (GAM, included in biomass pseudo-reactions) and non-growth-associated maintenance (NGAM, independent reaction) were respectively set to 29.89 and 2.2 mmol .g ^-1^.h^-1^ based on literature data for Chlamydomonas reinhardtii and *Nannochloropsis* sp. (Boyle and Morgan, 2009; Shah et al., 2017; Zhang et al., 2014).

### *In silico* predictions and validation

#### Validation of iMgadit23 quality with MEMOTE

Overall quality of iMgadit23 was evaluated using MEMOTE, a test suite for standardized quality assessment of GEMs (Lieven et al., 2020). The general properties of iMgadit23 were compared to the other GEMs of *Nannochloropsis* and *Microchloropsis*. General model features are summarized in Table 1 and complete MEMOTE reports are available in Supplemental Files S3-S6. iMgadit23 constitutes a high-quality model whose features include a multi-compartmentalized network encompassing most relevant microalgae compartments. Excluding reactions converting photons into chemical energy, all model reactions are charged and mass balanced. Only a low fraction of model reactions (10%, 99 reactions) can carry unlimited flux under model default medium (see Material and Method section). 27.3% of model reactions (636 reactions) lack a Gene Protein Reaction rule (GPR), most of them (319) being transport reactions or involved in lipid metabolism (282) due to gaps of knowledge of genes encoding lipid metabolism in *M. gaditana*. Among iMgadit23 genes, two pseudo-genes were introduced: (1) “Spontaneous”, associated to spontaneous reactions such as CO_2_, O_2_ or photon diffusion through membranes and (2) “NO_GENE”, associated to artificial reactions linked to modeling constraints (Biomass, NGAM, Exchange and Equivalence reactions). “OR” operator was used in GPRs to associate multiple genes to a common reaction. A total of 220 model metabolites are orphan (metabolites that are only consumed but not produced by reactions in the model) or dead-end (metabolites that can only be produced but not consumed by reactions in the model), resulting in a relatively high fraction of blocked reactions (19%, 443 reactions). Blocked reactions are generally caused by network gaps resulting from lack of metabolism knowledge. Nevertheless, in iMgadit23, blocked reactions do not prevent biomass production and are significantly fewer than in iNS937 and iRJ1321 where respectively 814 reactions (34.71%) and 930 reactions (48.49%) are blocked.

**Table 1.**
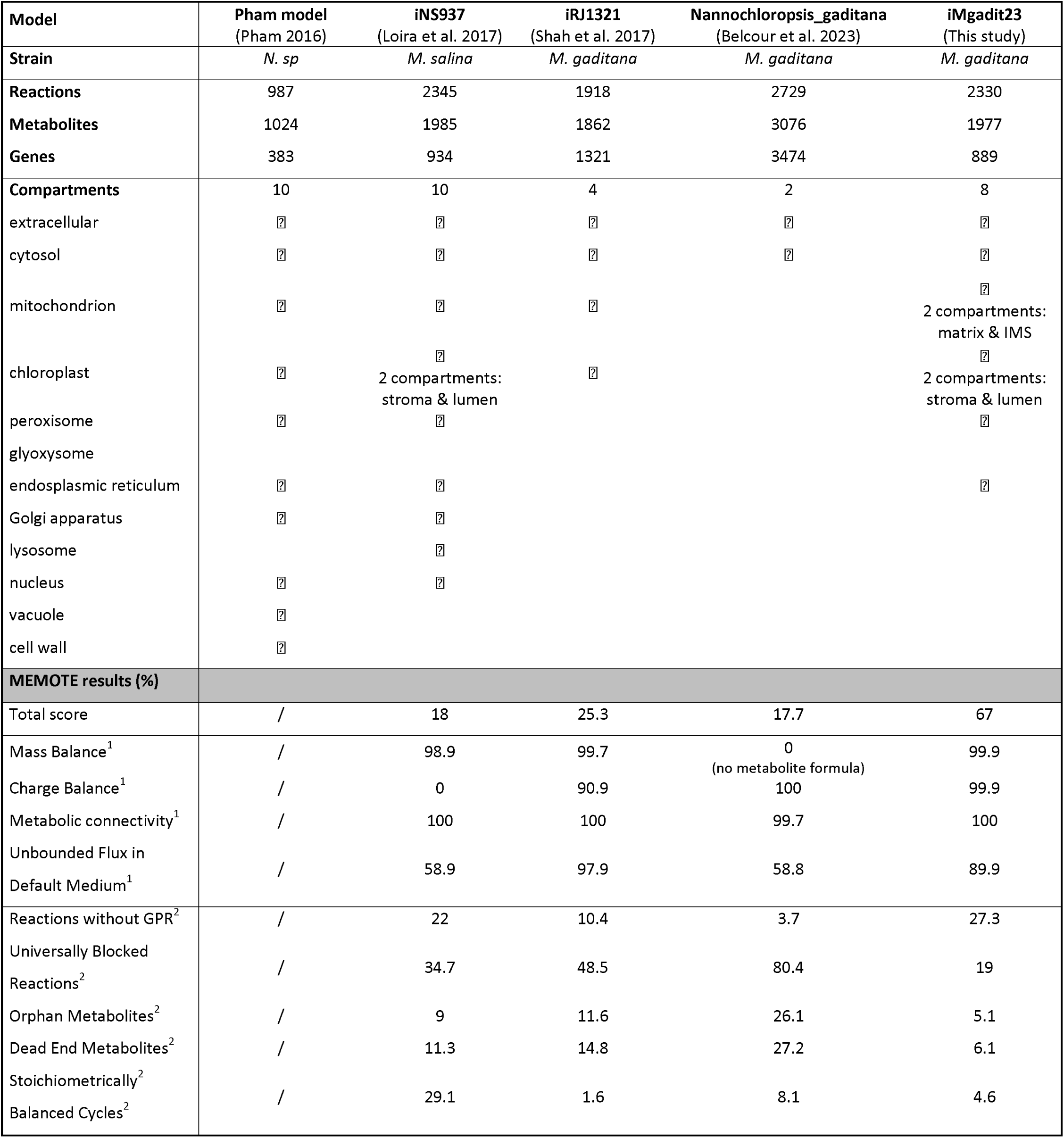
Properties of iMgadit23 and other *Nannochloropsis*, *Microchloropsis* GEM.

Detailed MEMOTE results also highlight high quality of lipid modeling in iMgadit23: among 1017 reactions describing lipid metabolism, only 61 (6%) are blocked. Moreover, some of these reactions are blocked because some lipids are not consumed in biomass reaction. When adding an artificial BOF reaction consuming all glycerolipids synthetized in model, number of lipid reactions blocked drops to 38. On the contrary, among 536 reactions representing lipid metabolism in iRJ321, 254 (47%) are blocked. Likewise, in iNS037, lipid metabolism is described by 827 reactions, 233 of which (28%) being blocked.

Finally, a “Total model score” of 67% was calculated by MEMOTE for iMgadit23. Both model consistency and annotation are considered to calculate this score. Consistency score of iMgadit23 is 55.7% because the model failed stoichiometry consistency test. All the different published models tested with MEMOTE in this study are also stoichiometrically inconsistent due to light energy conversion into chemical energy, breaking mass conservation law in models. When removing both photosystem reactions (PSII_h and PSI_h) from iMgadit23, the model becomes stoichiometrically consistent, leading to a consistency score of 98.5% and a total score of 79.6%, within the standard GEMs quality. Indeed, in 2020, a quality control assessment of the 108 SBML-GEMs of the BiGG database using MEMOTE revealed an average consistency and total scores of 92.18% and 72.54% (Norsigian et al., 2020)

### Quantitative validation

Three experimental maximum nitrate (NO_3_) uptake rates measured in *M. gaditana* cultivated under photoautotrophy (Rafay et al., 2020) were successively used to constrain NO_3_ uptake in iMgadit23 and pFBA was used to predict maximal growth rate. As presented in Figure 4A, model predicts an average maximum growth rate of 0.0253 ± 0.0014 h^-1^ which is slightly higher than maximum growth rate reported experimentally: 0.0183 ± 0.0038 h^-1^ (Rafay et al., 2020). Nevertheless, iMgadit23 predictions are in agreement with the maximal exponential growth rate values reported in literature for *M. gaditana* under photoautotrophic conditions with NO_3_ as nitrogen source, ranging from 0.01 to 0.044 h^-1^ with an average of 0.027 h^-1^ (Gentile and Blanch, 2001; Kareya et al., 2020; Kim et al., 2014; Ren and Ogden, 2014). iMgadit23 predictions are also consistent with a recently developed kinetic growth model of *M. gaditana*, predicting a maximal growth rate of 0.0256 h (Lima et al., 2022).

**Figure 4.**
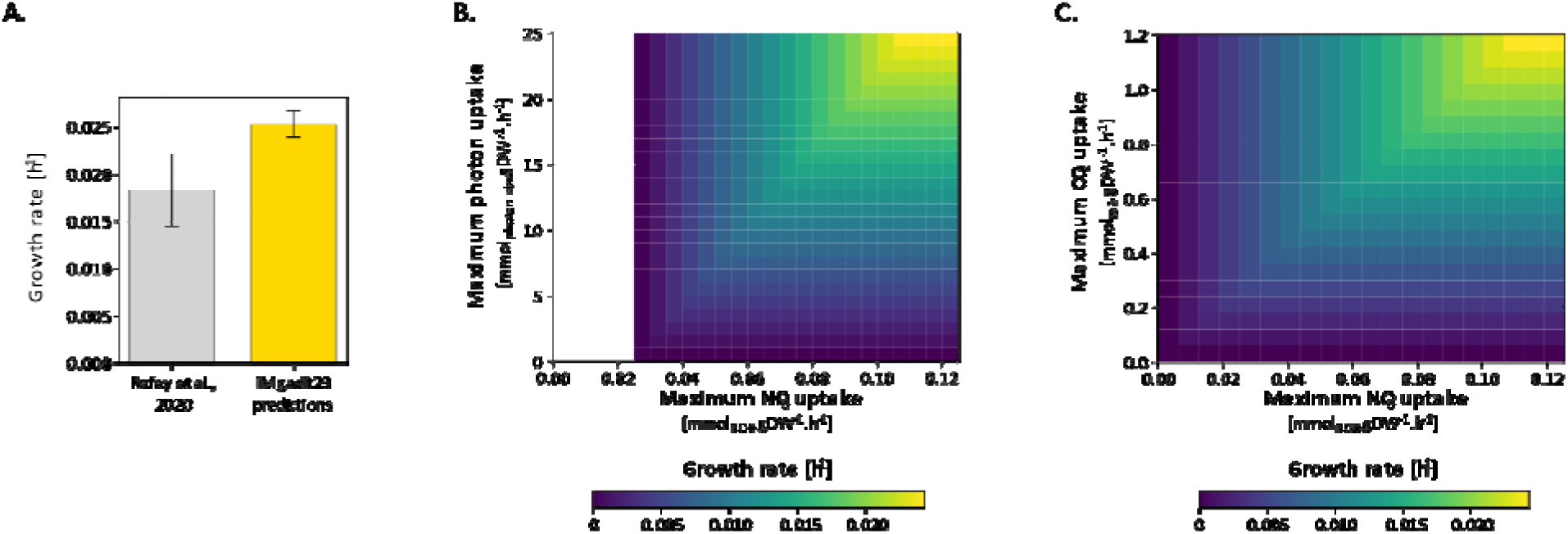
Predicted optimal growth rates of *M. gaditana* as a function of nitrogen, light and carbon dioxide. (A) NO_3_ uptake rates reported by (Rafay et al., 2020) were used to constrain iMgadit23. Maximum growth rates were predicted by pFBA. Phenotypic phase plane results provide a global view of how changes the optimal predicted growth depending in (B) of the NO_3_ and photon uptake and in (C) of the NO_3_ and CO_2_ uptake (color is used as third dimension to represent growth rate).

pFBA simulation results were used to calculate the ratio of oxygen evolved per carbon dioxide fixed. Ratio of 1.37 mol_O2_.mol_CO2_ predicted by iMgadit23 model is consistent with the experimental data reported for *Nannochloropsis* sp. under photoautotrophic conditions with NO_3_ as nitrogen source: ranging from 1.17 to 1.95 mol_O2_.mol_CO2_ (Zhang et al., 2014).

In microalgae, growth depends on numerous environmental parameters, notably availability of carbon, nitrogen sources and light intensity. Phenotypic phase plane analyses using pFBA approach were performed to provide an overview of how predicted optimal growth rate is impacted by changes in CO_2_, NO_3_ and photon availability (Figure 4B and C).

### Qualitative validation

*M. gaditana* as well as other *Nannochloropsis* and *Microchloropsis* species are able to grow in various conditions including: photoautotrophy with different nitrogen sources, heterotrophy and mixotrophy on different carbon sources (Fang et al., 2004; Gim et al., 2016; Kilian et al., 2011; Loira et al., 2017). iMgadit23 was tested for its ability to capture *M. gaditana* growth in 12 different conditions: heterotrophic growth on seven different carbon sources, mixotrophic growth on glucose and photoautotrophic growth on four nitrogen sources. Except for heterotrophic growth on ethanol, which is not predicted by iMgadit23 because ethanol metabolism is not represented in the model, all predictions meet experimental phenotypes (Table 2). In addition, growth phenotype of three knockout (KO) mutant strains was predicted in different media conditions. Model predictions are consistent with in vivo phenotypes for nitrate reductase KO (Naga_100699g1) and nitrite reductase KO (Naga_100852g1) tested for growth on different nitrogen sources and alternative oxidase KO (Naga_100568g3) (Bo et al., 2021; Kilian et al., 2011). Only the Bubblegum ACS MgACSBG (Naga_100014g59) is predicted as non-essential by iMgadit23, whereas (Billey et al., 2021) reported that this gene might be essential. *M. gaditana* genome encodes six annotated ACSs (Naga_100014g59, Naga_100012g66, Naga_101051g1, Naga_100047g8, Naga_100649g1 and Naga_100035g43) (Billey et al., 2021). Since specificities of ACS enzymes for fatty acids remain to be characterized, all ACSs genes were associated to cytosolic ACS reactions in iMgadit23 with the logical connector “OR”. Thus, inactivation of only one of ACS coding gene in iMgadit23 does not impact any ACS reaction functionality. Further analysis of ACS reactions is presented in the “Model use case” section.

**Table 2.** iMgadit23 growth predictions in different conditions.

| Condition | Medium | Gene KO | In vivo | In silico | TFPN | Species | Reference |
| --- | --- | --- | --- | --- | --- | --- | --- |
| Heterotrophy | Glucose |  | + | + | TP | *N. sp* | (Fang et al. 2004) |
|  | Ethanol |  | + | - | FN | *N. sp* | (Fang et al. 2004) |
|  | Xylose |  | + | + | TP | *N. oculata* | (Gim et al. 2016) |
|  | Rhamnose |  | + | + | TP | *N. oculata* | (Gim et al. 2016) |
|  | Fructose |  | + | + | TP | *N. oculata* | (Gim et al. 2016) |
|  | Sucrose |  | + | + | TP | *N. oculata* | (Gim et al. 2016) |
|  | Galactose |  | + | + | TP | *N. oculata* | (Gim et al. 2016) |
| Mixotrophy | Glucose |  | + | + | TP | *N. sp* | (Fang et al. 2004) |
| Photoautotrophy | Nitrite |  | + | + | TP | *N. sp* | (Kilian et al. 2011) |
|  | Nitrate |  | + | + | TP | *N. sp* | (Kilian et al. 2011) |
|  | Ammonium |  | + | + | TP | *N. sp* | (Kilian et al. 2011) |
|  | Urea levels |  | + | + | TP | *M. salina* | (Loira et al. 2017) |
| Knock-Out | Nitrite | Naga_100699g1 (Nitrate reductase) | + | + | TP | *N. sp* | (Kilian et al. 2011) |
| (Photoautotrophy) | Nitrate | Naga_100699g1 (Nitrate reductase) | - | - | TN | *N. sp* | (Kilian et al. 2011) |
|  | Ammonium | Naga_100699g1 (Nitrate reductase) | + | + | TP | *N. sp* | (Kilian et al. 2011) |
|  | Nitrite | Naga_100852g1 (Nitrite reductase) | - | - | TN | *N. sp* | (Kilian et al. 2011) |
|  | Nitrate | Naga_100852g1 (Nitrite reductase) | - | - | TN | *N. sp* | (Kilian et al. 2011) |
|  | Ammonium | Naga_100852g1 (Nitrite reductase) | + | + | TP | *N. sp* | (Kilian et al. 2011) |
|  | Nitrate | Naga_100568g3 (Alternative oxidase) | + | + | TP | *M. gaditana* | (Bo et al. 2021) |
|  | Nitrate | Naga_100014g59 (*MgACSBG*) | - | + | FP | *M. gaditana* | (Billey et al. 2021) |

Taking everything into account, iMgadit23 demonstrates good capacity for growth/no-growth predictions, with a sensitivity of 0.94, a specificity of 0.75 and an accuracy of 0.9. Good capacity of iMgadit23 to predict growth on different cultivation conditions both quantitatively and qualitatively highlights its usefulness to evaluate the biological capabilities of *M. gaditana* on different conditions.

### *Model use case*: characterization of *acsbg* mutant

One of the strengths of GEMs is to be a support for researchers working on metabolism or molecular engineering. For this reason, a mutant strain described in the literature was chosen to evaluate iMgadit23. The MgACSBG gene was studied by Billey and co-authors (Billey et al., 2021). In their work, they report that despite numerous attempts, they were unable to obtain a MgACSBG knockout, suggesting that reactions catalyzed by the MgACSBG protein may be vital for *M. gaditana*. As previously mentioned, iMgadit23 failed to predict MgACSBG essentiality due to lack of specificity information regarding ACS enzymes.

Billey and co-authors generated point mutations in MgACSBG gene and characterized its impact on lipid profile. MgACSBG#31 mutant from (Billey et al., 2021) (referred to as “acsbg mutant” in our study) harbors a local mutation in MgACSBG leading to three amino acid changes in the MgACSBG protein. Structural modeling of the mutant protein suggests that accessibility of FAs to the catalytic site is affected. Glycerolipid profile is highly impacted by the mutation. Mutant strain notably showed (1) an increase in DGTS, (2) a very large increase in TAGs, (3) an overall decrease of 16:1 in glycerolipids, (4) a specific increase of unsaturated 18-carbon acyls in PC and (5) a decrease of 20-carbon acyls in DGTS. Altogether, results presented by (Billey et al., 2021) suggest a role of MgACSBG in the production of 16:1-CoA and 18:3-CoA. Detailed analysis of lipid composition being provided in the publication, this gene makes a good candidate for iMgadit23 evaluation.

### ACS reactions deletion

First, essentiality analysis of each cytosolic ACS reaction of iMgadit23 was conducted to determine which reactions, when deleted one by one in silico, significantly reduces or eliminates biomass production. Nomenclature of the ACS reactions uses the fatty acid one: for example, ACS140_c corresponds to the reaction of thio-esterification of fatty acid 14:0 to CoA occurring in the cytosol. Most reactions (ACS140_c, ACS160_c, ACS161_c, ACS181_c, ACS183_c, ACS204_c and ACS205_c) are predicted as essential by iMgadit23, whereas deletion of ACS120_c, ACS180_c, ACS182_c and ACS203_c do not affect predicted growth (Supplemental File S7). These results suggest that MgACSBG might be specific for one or more of the FAs: 14:0, 16:0, 16:1, 18:1, 18:3, 20:4 or 20:5.

### Differential flux analyses

In a second step, effect of the MgACSBG point mutation was modelled by introducing a new biomass reaction in iMgadit23 based on acsbg biomass composition (specifically lipid composition) and by constraining this acsbg strain-specific biomass reaction to the experimental acsbg growth rate (Billey et al., 2021). Flux simulations were then performed to identify which reaction could lead to the observed changes in acsbg growth rate and biomass composition compared to WT strain, and thus infer the specific metabolic role and location of the targeted MgACSBG.

At first, flux distributions were predicted by pFBA for WT and acsbg strains. For each simulation, appropriate BOF was constrained to experimental growth rate (0.030 h^-1^ for WT, 0.015 h^-1^ for acsbg) and pFBA was conducted with minimization of photon uptake rate as objective function. To facilitate reading, these flux distributions will be respectively termed pFBA_WT and pFBA_acsbg. For each model reaction, log2(pFBA_acsbg/pFBA_WT) was calculated (Figure 5A). A positive log2 fold change (log2(FC)) indicates that reaction flux is increased in acsbg compared to WT. On the contrary, a negative log2(FC) indicates a decrease of the reaction flux in the mutant. Reactions with −0.26 ≤ log2(FC) ≤ 0.26, corresponding to a flux change lower than 20%, were considered as non-significantly impacted (See Material and Methods section for more details).

**Figure 5.**
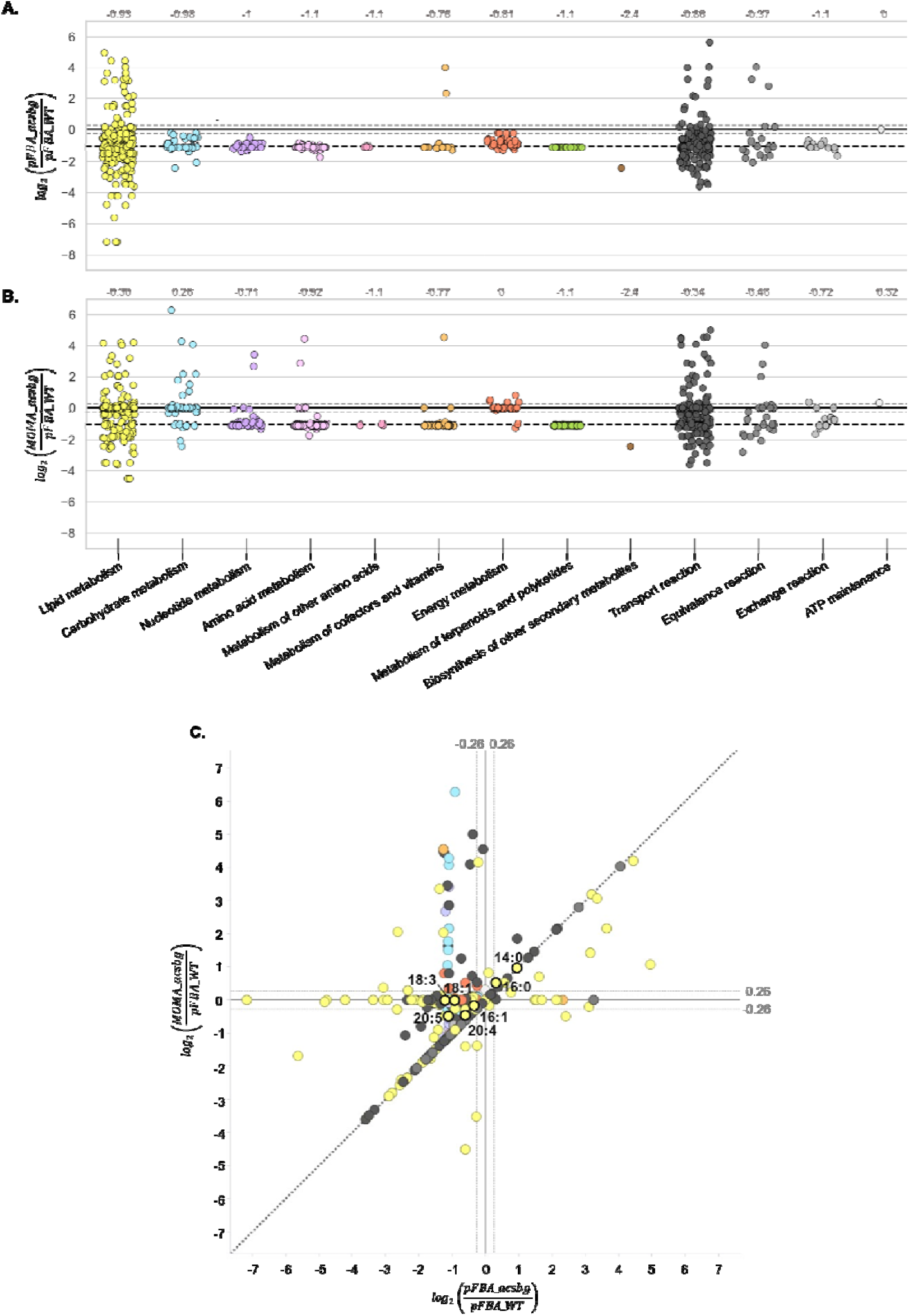
Results of the differential flux analyses between WT and acsbg strains. Comparison of flux values for both strains for each model reaction grouped by subsystem predicted with (A) pFBA approach (log2(pFBA_acsbg/pFBA_WT)) and (B) MOMA approach (log2(MOMA_acsbg/pFBA_WT)). A positive log2(FC) indicates an upregulated reaction in mutant strain whereas a negative value indicates a downregulated reaction. Average log2(FC) of the subsystem is indicated in grey. Dashed black line indicates log2(µ_acsbg_/µ_WT_) = −1, dashed grey lines indicate significant thresholds of −0.26 and 0.26 corresponding to an up/downregulation of 20% in acsbg compared to WT. (C) Comparison of log2(FC) between both pFBA and MOMA approaches. Diagonal line represents identical predictions with both methods. Reactions predicted to be upregulated in acsbg by both methods are in the upper right corner, reactions predicted to be downregulated by both methods are in the bottom left corner. Reactions predicted to be downregulated in pFBA_acsbg but upregulated in MOMA_acsbg are in the upper left corner. Reactions predicted to be upregulated in pFBA_acsbg but downregulated in MOMA_acsbg are in the bottom right corner. Dashed grey lines indicate significant threshold of −0.26 and 0.26. Cytosolic ACS reactions are highlighted in black.

Most reactions are downregulated in pFBA_acsbg compared to pFBA_WT (Figure 5A). In total, among 770 reactions predicted to have a flux in pFBA_WT or pFBA_acsbg, 606 have a significantly lower flux in mutant compared to WT. This downregulation tendency can be explained by experimental strains growth rates (µ_WT_ = 0.030 h^-1^, µ = 0.015 h^-1^) being applied as simulation constraints. Thus, many reactions are predicted to have a log2(pFBA_acsbg/pFBA_WT) close to log2(µ_acsbg_/µ_WT_) = −1. Despite these simulation constraints, 50 reactions, mostly involved in lipid metabolism and lipid transport, are predicted to be upregulated in pFBA_acsbg compared to pFBA_WT. Lipid metabolism and lipid transport are the most impacted subsystems in terms of number of up and downregulated reactions but also in terms of log2(FC) ranges.

Then, we applied minimization of metabolic adjustments (MOMA), another popular COBRA method developed to find a flux distribution for a perturbed condition by minimizing the Euclidean distance from a reference flux distribution (Segrè et al., 2002). MOMA is considered as more suitable to predict flux distributions in mutant organisms. acsbg BOF was constrained to its experimental value, and fluxes were predicted with MOMA using pFBA_WT as reference distribution. Predicted fluxes will be referred to as MOMA_acsbg. log2(MOMA_acsbg/pFBA_WT) was calculated for each iMgadit23 reaction (Figure 5B). Only flux changes higher than 20% were considered significant. Among the 1009 reactions predicted to have a flux in pFBA_WT or MOMA_acsbg, 373 (including 79 reactions of lipid metabolism) are significantly downregulated in acsbg, 80 (including 26 reactions of lipid metabolism) are significantly upregulated and 253 reactions have similar fluxes for both strains.

pFBA and MOMA predictions for acsbg were then compared to identify reactions predicted to be up- or down-regulated in mutant by both approaches, with a focus on cytosolic ACS reactions (Figure 5C). 31 reactions are upregulated in pFBA_acsbg and MOMA_acsbg compared to pFBA_WT, with 18 of these reactions involved in lipid metabolism (Figure 5C). Interestingly, ACS140_c and ACS160_c are part of these significantly upregulated reactions. On the other hand, 366 reactions, including 74 reactions of lipid metabolism notably ACS204_c and ACS205_c, are downregulated both in pFBA_acsbg and MOMA_acsbg. ACS161_c is significantly downregulated in pFBA_acsbg and non-significantly downregulated in MOMA_acsbg with log2(MOMA_acsbg/pFBA_WT) = −0.17 (corresponding to flux decrease of 12%). For ACS181_c and ACS183_c, fluxes are significantly decreased in pFBA_acsbg, but log2(MOMA_acsbg/pFBA_WT) = 0. For both WT and acsbg, a null flux is predicted for ACS120_c, ACS180_c, ACS182_c and ACS203_c by pFBA and MOMA. All reactions significantly predicted as up regulated or downregulated in acsbg by both methods are detailed in Supplemental File S8. pFBA_WT, pFBA_acsbg, MOMA_acsbg and log2(FC) for each model reaction are detailed in Supplemental File S9. The lipid metabolism results are also detailed in Figure 6.

**Figure 6.**
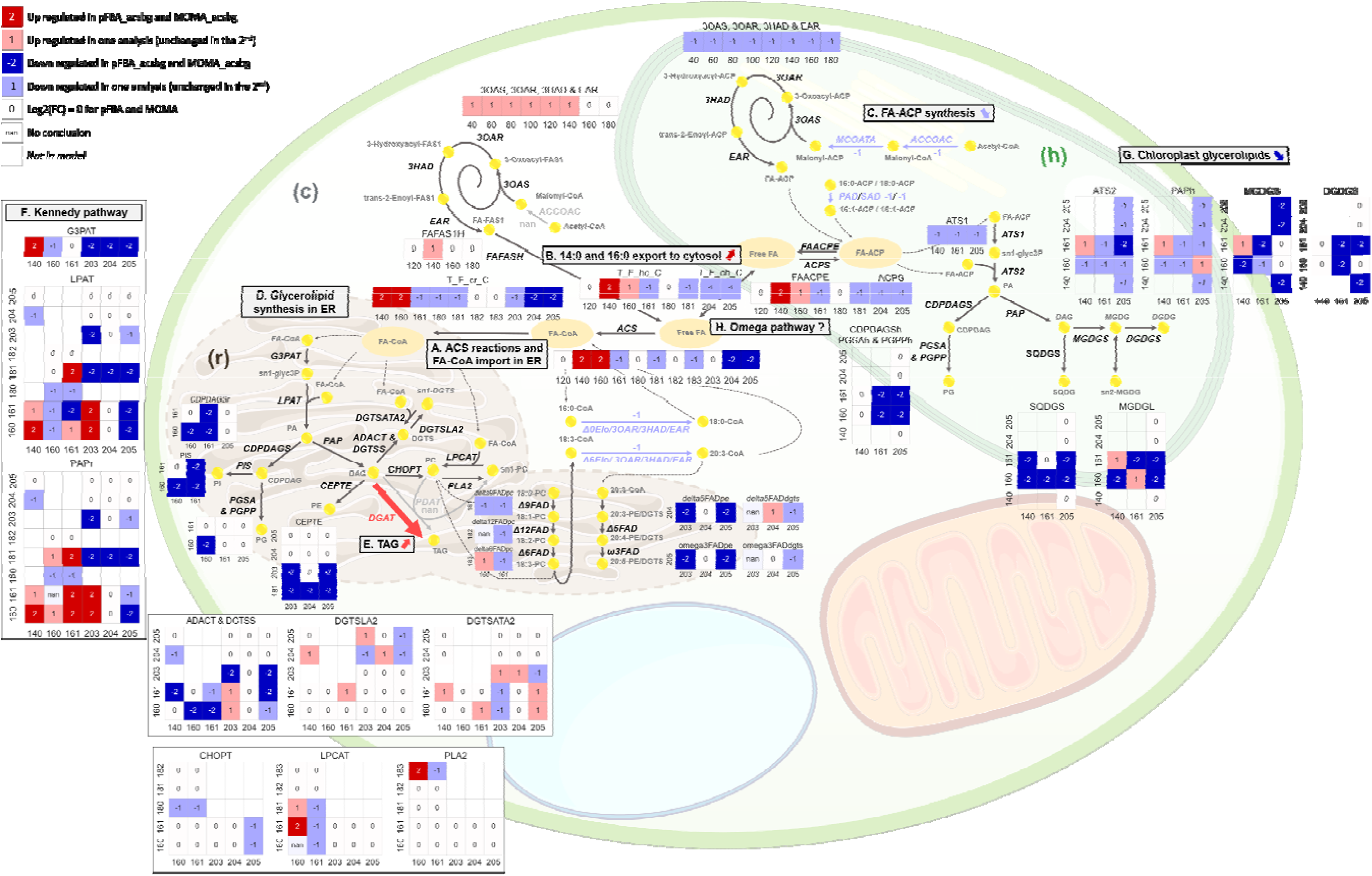
pFBA and MOMA predictions for lipid metabolism. For each reaction results are shown for all acyl chains couples: sn-1 (abscissa-axis) and sn-2 (ordinate axis). Red colors indicate reactions significantly upregulated in acsbg with both methods (dark red) or one method (light red). Blue colors indicate reactions significantly downregulated in acsbg with both methods (dark blue) or one method (light blue). Reactions for which same fluxes for WT and acsbg are predicted by both methods are indicated by 0. Reactions predicted up by one method but down by second method are notified as “nan” to indicate absence of conclusion. Finally, empty value indicates that reaction does not exist in iMgadit23. Main pathways of lipid metabolism are named and labelled from A to H. Cytosol, chloroplast and ER are indicated as (c), (h) and (r).

Constraint-based modeling leaves open the possibility of alternate optimal solutions meaning that the same objective value can be achieved by a diverse set of flux distributions. Flux variability analysis (FVA) determines the possible range of flux values (i.e. minimum and maximum values for each reaction) which is allowable with the given objective value (Gudmundsson and Thiele, 2010). To give an overview of flux variability within ACS reactions, we performed flux variability analyses (FVA) to determine the range of possible reaction fluxes that still satisfy the original optimal objective value predicted by pFBA (Table 3). FVA approach, with 100% optimality constraint, was applied for WT and acsbg strains by constraining appropriate BOF to experimental growth rate, with minimization of photon uptake as objective function. For each reaction, the fractional overlap in flux range between WT and acsbg was calculated as defined in Equation 5 (See Material and Methods). Interestingly, no overlap is predicted for ACS140_c and ACS160_c confirming their upregulation in acsbg. No overlap is also predicted for ACS161_c, ACS181_c, ACS183_c and ACS205_c reinforcing pFBA_acsbg downregulation hypothesis. ACS204_c has a fractional overlap of 0.34, suggesting that its predicted upregulation in pFBA_acsbg can be due to variability of lipid metabolism.

**Table 3.** FVA results for cytosolic ACS reactions.

| Reaction ID | WT<br>[min,max]<br>($\mu\text{mol.gDW}^{-1}.\text{h}^{-1}$) | Acsbg<br>[min,max]<br>($\mu\text{mol.gDW}^{-1}.\text{h}^{-1}$) | Fractional overlap |
| --- | --- | --- | --- |
| ACS120_c | [0,0] | [0,0] | 1 |
| ACS140_c | [0.21,0.21] | [0.41,0.41] | 0 |
| ACS160_c | [3.1,4.3] | [4.9,5.3] | 0 |
| ACS161_c | [3.9,4.0] | [3.1,3.1] | 0 |
| ACS180_c | [0,1.2] | [0,0.43] | 0.36 |
| ACS181_c | [4.5,4.5] | [2.4,2.4] | 0 |
| ACS182_c | [0,0] | [0,0] | 1 |
| ACS183_c | [5.1,5.1] | [2.2,2.2] | 0 |
| ACS203_c | [0,0] | [0,0] | 1 |
| ACS204_c | [0.03,0.38] | [0.02,0.15] | 0.34 |
| ACS205_c | [0.4,0.9] | [0.2,0.4] | 0.07 |

To sum up regarding ACS reactions, all three approaches pFBA, MOMA and FVA agree in predicting an upregulation of thioesterification to CoA of long saturated FA chains 14:0 and 16:0, while a downregulation is observed for very long and unsaturated FA 20:5. Thio-esterification to CoA of FA 16:1 is also predicted to be significantly downregulated by pFBA and FVA, while MOMA approach only suggests a non-significant decrease (12% decrease in acsbg flux compared to WT). Two of the three approaches (pFBA and FVA) predict a significant downregulation of ACS 18:3 in acsbg. Finally, pFBA and MOMA results also predict downregulation of FA 20:4 thio-esterification to CoA. Nevertheless, FVA approaches predicts that WT and acsbg flux ranges for ACS 20:4 are overlapping (ξ = 0.34) suggesting that downregulation predicted by pFBA and MOMA for acsbg could arise from lipid metabolism variability within iMgadit23. All together, these results are consistent with the reported specificity of ACS proteins of the bubblegum family in the activation of long chains FAs and VLC-FAs with chain lengths from 16 to at least 24 carbons from studies in drosophila, rat and human (Min and Benzer, 1999; Song et al., 2007; Steinberg et al., 2000). Moreover, these results are in close agreement with the findings reported in (Billey et al., 2021), where a significant accumulation of 16:0 acyl-CoA, a significant depletion of 16:1 and 18:3 acyl-CoA, and a minor depletion of 20:5 acyl-CoA were observed in the acsbg mutant. It is important to keep in mind that only glycerolipidome (therefore excluding acyl-CoA profile) of acsbg mutant was used to add an ascbg mutant specifiC BOF in iMgadit23 and that no additional constraint was applied on ACS coding genes or reactions in model while conducing pFBA, MOMA and FVA. Still, ACS reactions are predicted to be impacted in acsbg simulations with all three approaches.

To go further in deciphering acsbg reorganization of lipid metabolism, pFBA and MOMA predictions for the full lipid metabolism were compared (Figure 6). Respective pFBA and MOMA log2(FC) values are also recapitulated in Supplemental Results, Tables S1 and S2 and Figure S2. As previously mentioned, cytosolic thio-esterification of free FA to CoA by ACS is predicted to be upregulated for 14:0, 16:0 FAs and downregulated for 16:1 (significant only in pFBA_acsbg), 20:4 and 20:5 FAs in acsbg. Similar results are predicted for FA-CoA transport to ER (Figure 6A). Plastidial 14:0-ACP and 16:0-ACP conversion to free FA (FAACPE reactions), followed by free FAs 14:0 and 16:0 export from chloroplast to cytosol (T_F_hc_C reactions) are also predicted to be upregulated in pFBA_acsbg (for both FAs 14:0 and 16:0) and MOMA_acsbg (only for 14:0) compared to pFBA_WT (Figure 6B). Regarding, FAACPE160_c and T_F_hc_C160, fluxes are significantly increased in pFBA_acsbg, but predicted increase is not significant with MOMA approach. Interestingly, while in pFBA_acsbg and MOMA_acsbg more 14:0-ACP and 16:0-ACP are converted to free FAs which are then exported from chloroplast, de novo synthesis of 14:0-ACP and 16:0-ACP is not predicted as upregulated (nor for any FA-ACP) in acsbg (Figure 6C). In pFBA_acsbg, FA-ACP biosynthesis is even predicted to be downregulated.

Impact of acsbg lipid profile on glycerolipid synthesis in ER is highly variable depending on sn-1, sn-2 acyl chains (Figure 6D). However, TAGs biosynthesis via DGAT reactions is widely predicted to be upregulated both in pFBA_acsbg and MOMA_acsbg (Figure 6E and Supplemental File S10). Additionally, many LPAT and PAP reactions, involved in TAGs biosynthesis via the Kennedy pathway, are predicted to be upregulated in acsbg, notably reactions incorporating 16:0 at the sn-2 position of ER lipids and reactions incorporating 16:1 both at the sn-1 and sn-2 positions of ER lipids (Figure 6F). Finally, an overall downregulation tendency of plastid glycerolipid synthesis is predicted in acsbg mutant with both methods (Figure 6G). These results are consistent with proposed role for MgACSBG (Billey et al., 2021), highlighting that iMgadit23 constitutes a valuable tool to decipher mutant phenotype, notably regarding lipid metabolism. pFBA, MOMA and FVA results for all model reactions are available in Supplemental File S9.

### Discussion and perspectives

#### iMgadit23 model

GEMs are powerful tools to explore microorganisms metabolism. In this work, iMgadit23 model of *M. gaditana* was reconstructed and manually curated. Model ability to qualitatively and quantitatively predict in vivo growth phenotypes under diverse environmental and genetic conditions was validated. Our results highlight value of iMgadit23 in guiding model-driven strain design and medium optimization to enhance growth or yields of target compounds.

A comprehensive 2D-pathway map of iMgadit23 content is available in JSON format (https://github.com/Total-RD/Mgaditana-GEM). This map can serve as a platform to visualize and analyze flux distributions previously predicted using COBRA methods. The map can also facilitate the exploration of other relevant biological datasets including metabolic fluxes, lipidomics, metabolomics, and transcriptomics data using the Escher tool (King et al., 2015). In addition, the map, together with iMgadit23 GEM, enables the execution of interactive FBA simulations through Escher-FBA, a web-based open-source application (Rowe et al., 2018). Within this tool, users can interactively and easily adjust various FBA simulation parameters, including flux bounds, objective functions, and reaction knockouts and immediately visualize the impact on simulation results without programming skills.

#### Photosynthesis modeling

In iMgadit23, light phase of photosynthesis is modeled by five reactions describing the LEF and the ATP synthase. Absence of CEF description was bypassed by the addition of PSQUANTUM_h reaction, introducing distinction between absorbed and LEF photons. This addition decreases the absorbed photon/ATP ratio from 8/3 to 10/3, leading to a predicted photosynthesis efficiency more consistent with experimental data. Conversion of 10 absorbed photons to 8 LEF photons was hypothesized based on literature, but this 10/8 ratio can be easily adjusted to modify absorbed photon/ATP ratio. Indeed, photosynthetic organisms have evolved alternative electron flows (AEF), such as CEF, allowing H^+^ translocation without producing NADPH, consequently impacting photon/ATP ratio. All AEFs have a distinct H^+^/e^-^ ratio, and their relative occurrences are constantly adapted to environmental conditions (CO_2_, O_2_, light quantity and quality, redox state, nutrient availability etc.) (Curien et al., 2016). Addition of PSQUANTUM pseudo-reaction constitutes a first step to better represent photon requirement of ATP production. One perspective to improve iMgadit23, would be to add AEF, starting with CEF. Impact of environmental conditions on flux distributions within these pathways as well as impact on photosynthesis yields would be interesting results to further characterize photoautotrophy in *M. gaditana*.

#### Modeling of the lipid metabolism

Quality and details of lipid metabolism modeling and BOF lipid composition highly impact GEMs flux predictions (Correa et al., 2020; Levering et al., 2016; Sánchez et al., 2019). Due to diversity of existing lipid entities, lipid metabolism tends to be the most complex part of metabolic models. One approach to get around lipid metabolism complexity, consists in considering that for each lipid metabolite sn-1, sn-2 (and sn-3 for TAGs) acyl chains are identical. This approach was used in iRJ1321 model, greatly simplifying lipid metabolism (Shah et al., 2017). Using the example of TAG biosynthesis, which is of special interest for biotechnology applications, ten acyl chains (14:0, 16:0, 16:1, 18:0, 18:1, 18:2, 18:3_ω3, 18:3_ω6, 20:4 and 20:5) are considered in TAG metabolites in iRJ1321. Instead of considering all possible arrangements with repetition of three among ten acyl chains (resulting in 10^3^ possible TAGs), only ten distinct TAGs with three identical acyl chains are described in iRJ1321 (tag140_c corresponding to TAG 14:0/14:0/14:0, tag160_c for TAG 16:0/16:0/16:0 etc.). In iMgadit23, eight acyl chains were considered for TAGs (14:0, 16:0, 16:1, 18:0, 18:1, 20:3, 20:4 and 20:5). Also, rather than considering only eight TAGs or describing all 512 (8^3^) putative possible TAGs, experimental data were used to determine existing TAGs (35 in total) in *M. gaditana* (Billey et al., 2021; Jouhet et al., 2017). Only reactions required for the metabolism of these 35 TAGs were added to model. For lipid synthesis in iMgadit23, unsaturation position within acyl chains was not taken into account. Consideration of unsaturation positions, notably regarding VLC-PUFAs could constitute an improvement of lipid metabolism description in our model.

In GEMs, the level of detail of lipid metabolism is often interrelated with the BOF lipid composition which depends on availability of detailed data. Indeed, exhaustive quantification of all lipid entities is not often available, resulting in BOF stoichiometric coefficients calculated from approximated lipid data. Here, data used for BOF lipid composition rely on on FA quantification by gas chromatography (GC) and glycerolipid quantification by high-pressure liquid chromatography–tandem mass spectrometry (HPLC-MS/MS). Combination of both methods resulting in the precise quantification of the fatty acids present in each glycerolipid class (Billey et al., 2021; Jouhet et al., 2017). The strength of iMgadit23 is to be built and validated on such detailed lipidomic data that makes this model available for a deep analysis of lipid metabolism.

Finally, in terms of reaction description, iMgadit23 constitutes an up-to-date overview of knowledge of *M. gaditana* lipid metabolism. Prediction of flux distribution together with model quality assessment with MEMOTE, revealed that among the different lipid pathways, all are functional except steroid biosynthesis. Due to gaps of knowledge of sterol metabolism, its curation in our model would need further investigation. Sphingolipid metabolism is missing in iMgadit23. This pathway is described in both iNS937 and iRJ321 but non-functional, most reactions being blocked, likely due to the limited knowledge of sphingolipid metabolism in *Nannochloropsis* and *Microchloropsis*. Integrating functional sphingolipid metabolism into iMgadit23 would also require further biochemical and genetic characterizations. Gaps of knowledge for microalgal lipid synthesis, editing and degradation pathways still need to be filled notably for strains containing plastids with four limited membranes such as *M. gaditana* (Hoffmann and Shachar-Hill, 2023; Petroutsos et al., 2014). As a result, in iMgadit23, some reactions were added, although their existence in *M. gaditana* awaits experimental confirmation. For example, 18:0 and 18:1 synthesis in chloroplast has been described in higher plants but no clear evidence of its existence in microalgae has been reported so far. Similarly, acyl editing on PC, DGTS and MGDG is presumed to exist in *M. gaditana* although its actual existence requires validation (Hoffmann and Shachar-Hill, 2023). Lipid metabolism characterization of *M. gaditana* and microalgae, in general, is an ongoing research topic. Continuously updating iMgadit23 with new knowledge could provide valuable insights into its quality and predictive capabilities. Some published microalgal GEMs already tend to detail pathways associated with lipid biosynthesis and metabolism. However, in (Correa et al., 2020), Correa et al. reported that of 45 microalgal GEMs examined, only 26 included more than 50 reactions related to lipids. These 45 models included an average of 348 reactions describing lipid metabolism while in iMgadit23 lipid metabolism is described by 1017 reactions.

Due to the precision in lipid metabolism modeling, including reaction description and compartmentation, and the high-quality data used for calculating the lipid composition of the BOF, iMgadit23 stands as one of the most comprehensive models for lipid modeling in microalgae.

#### Analysis of acsbg mutant phenotype

Given the complexity and variability of lipid metabolism, characterizing and interpreting mutant strain phenotypes is very challenging, often requiring lots of experimental investigation that are costly and time consuming. In this study, we employed iMgadit23 to elucidate the acsbg mutant phenotype. Two COBRA methods, pFBA and MOMA, were applied to predict impact of acsbg mutant lipid profile on reaction fluxes. The pFBA approach finds an optimal flux distribution which maximizes the objective function and minimizes the total sum of the absolute value of fluxes. On the other hand, MOMA minimizes the Euclidean distance from a reference flux distribution. Despite the difference in their mathematical objective, overall predictions for acsbg with pFBA and MOMA approaches consistently and effectively predicted the acsbg metabolic phenotype. In total 470 reactions are robustly predicted as up or downregulated by both methods. They notably predicted (1) the upregulation of cytosolic ACS 14:0 and 16:0 reactions; (2) the downregulation of ACS 20:5, 16:1 reactions (ACS161_c downregulation being significant only in pFBA_acsbg); (3) the upregulation of many reactions of TAGs biosynthesis via the Kennedy pathway, notably reactions incorporating 16:0 and 16:1 at the sn-2 position of ER lipids; and (4) an increase export of free FAs 14:0 and 16:0 from chloroplast without increasing de novo FA-ACP synthesis. These predictions are consistent and partially confirm without any a priori the hypotheses presented by (Billey et al., 2021), particularly the proposed role for MgACSBG in the production of 16:1-CoA in the cytosol, prior to 16:1 incorporation at the sn-2 position of ER lipids via the Kennedy pathway. One important assumption for this proposed model is that positive feedback is exerted on the plastid, exporting more FAs. This assumption was validated by iMgadit23 for 14:0 and 16:0 FAs. Nevertheless, a second role of MgACSBG in the production of 18:3-CoA was also proposed by (Billey et al., 2021) mainly based on their in vivo analysis of acyl-CoA profiles. This analysis revealed a significant depletion in 16:1-CoA and 18:3-CoA along with an accumulation of 16:0-CoA and 18:1-CoA in acsbg mutant. While pFBA predict a significant downregulation of ACS183_c in acsbg (confirmed by FVA) with a log2(pFBA_acsbg/pFBA_WT) = −1.2, which is lower than log2(µ_acsbg_/µ_WT_) = −1; MOMA does not predict this downregulation (log2(MOMA_acsbg/pFBA_WT) = 0). This may be partially explained by the steady state assumption inherent in COBRA methods used in this study. Accumulation or depletion of these metabolites could not be taken into account in the differential flux analyses, resulting in the absence of ACS183_c downregulation in MOMA_acsbg.

#### pFBA approach

Regarding pFBA approach, on the one hand using experimental growth rates (µ_WT_ = 0.030 h^-1^, µ = 0.015 h^-1^) to constrain BOF enables consideration of the mutant growth defect in predicted flux distributions. On the other hand, this approach has the drawback of potentially masking downregulated reactions.

#### MOMA approach

Numerous energy-consuming reactions, such as NGAM, alternative oxidase (AOX) along with other futile cycles are activated in MOMA_acsbg (these reactions have a flux in acsbg mutant but not in WT). Cytosolic FA 14:0 synthesis is activated in MOMA_acsbg while 14:0 degradation is also activated. All these activations may be attributed to similar fluxes in photosynthesis, but with BOF demand being only half as much in acsbg compared to WT. Consequently, the excess energy not utilized for biomass production must be consumed. Interestingly, peroxisomal β-oxidation of FAs 20:5, 20:4, 20:3, 18:3, 16:1 and 14:0 was predicted to have a significant flux in MOMA_acsbg. This activation of peroxisomal FAs degradation comes with the activation of glyoxylate cycle and photorespiration, as well as increased fluxes in TCA cycle, citrate/malate/oxaloacetate metabolism, serine/glycine metabolism and pentose phosphate pathway. In silico, degradation of 20:5, 20:4, 20:3, 18:3, 16:1 and 14:0 FAs suggests that these FAs are less needed for incorporation into glycerolipids in MOMA_acsbg. MOMA, along with FBA and pFBA approaches, relies on steady state assumption i.e. total the total production of a metabolite must balance with its total consumption. Consequently, these methods cannot predict the accumulation of FAs, resulting in their degradation in MOMA_acsbg. In vivo, the pool of available FA-CoAs is modified (Billey et al. 2021). Activation and upregulation of β-oxidation, glyoxylate cycle, photorespiration and TCA cycle is therefore a biologically plausible assumption that need to be experimentally validated.

In silico differential flux analyses between WT and acsbg strains demonstrate the utility of iMgadit23 as a valuable tool to decipher mutant phenotypes and ultimately increase our understanding of *M. gaditana* metabolism. Similar methodologies, relying on context-specific biomass objective functions, could be applied to elucidate the metabolic phenotype resulting from metabolic engineering or variation in culture parameters including light, nitrogen or phosphate availability.

## Conclusion

GEMs are powerful tools to explore the metabolism of biological systems. In this study, we reconstructed, manually curated and experimentally validated iMgadit23, a high-quality GEM for the industrially promising oleaginous microalga *M. gaditana*, with a special focus on lipid metabolism. To date, iMgadit23 is the most comprehensive GEM available for *Nannochloropsis* and *Microchloropsis* species, especially with respect to lipid metabolism. Our analyses showed that iMgadit23 can accurately predict growth phenotypes under different media and genetic conditions, both qualitatively and quantitatively. Moreover, the in silico analysis of the acsbg mutant phenotype highlights the potential of iMgadit23 to elucidate the physiological behavior and metabolic states of *M. gaditana* in response to genetic or environmental changes. We believe that iMgadit23, together with its comprehensive 2D-pathway map, will be valuable tools for identifying novel metabolic engineering strategies aimed at enhancing lipid synthesis and accumulation in *M. gaditana*, thus opening new opportunities using this strain as a platform for various industrial applications. This approach might enable in the future the optimization of cultivation processes and the design of new strains with enhanced lipid production.

## Material and Methods

### Draft model automatic reconstruction and manual curation

iMgadit23 was reconstructed based on the complete genome sequence of *M. gaditana* B-31 (GenBank accession number GCA_000569095.1) (Corteggiani Carpinelli et al., 2014), according to the methodology presented in (Pereira et al., 2018). Briefly, model reconstruction was carried out in three different phases: (1) genome reannotation, (2) draft metabolic model building, (3) model completion and refinement by network gaps filling, compartmentalization of reactions and metabolites, addition of missing pathways and correction of inconsistencies based on literature and expert knowledge.

1. The functional annotation of genes was based on homology searching methods like BLAST (Altschul et al., 1990) and HMMER (Eddy, 1998) against protein sequence databases, such as UniProt (UniProt Consortium, 2018) to find the best aligned sequences. The best matches were manually revised and the corresponding Enzyme Commission (EC) number(s) was associated to each gene. Functional scores were also associated providing evidence for the assigned metabolic function(s), with weighed scores ranging between 0 and 1 (1 corresponding to a high confidence score) attributed by the curator after examination of homology results and previously reported knowledge.
2. The association of biochemical reactions is carried out based on the associated EC number(s), and once again manually revised to guarantee the compilation of strain-specific metabolic features. Spontaneous and transport reactions from databases like KEGG (Kanehisa and Goto, 2000) and Transporter Classification Database (TCDB) (Saier et al., 2014) are also added. Some metabolic reactions, although not associated with genes, were also included due to evidence found in literature.
3. Model refinements include several steps from gap filling to compartmentalization of reactions that allow to reach a high-quality GEM with predictive value and strain-specific information. More details on these steps can be found in (Pereira et al., 2018). One of the most relevant steps is the definition of a biomass reaction representing the basic macromolecular composition of *M. gaditana* using strain-specific data in terms of macromolecular contents (PROTEINS, DNA, RNA, LIPIDS, CARBOHYDRATES and PIGMENTS). Pseudo-reactions representing the biosynthesis of glycerolipid classes (TAG, DAG, PC, PE, PI, PG, DGTS, DQDG, MGDG and DGDG) were also included in the GEM.

### Biomass objective functions composition

iMgadit23 contains two independent BOFs, representing respectively biomass composition of WT and MgACSBG#31 mutant strains (Billey et al., 2021). Each biomass objective function involves 17 pseudo-reactions and pseudo-metabolites representing BIOMASS, the different macromolecules (DNA, RNA, PROTEINS, CARBOHYDRATES, PIGMENTS and LIPIDS) and glycerolipid classes (DAG, TAG, PC, PE, PI, PG, DGTS, DGDG, MGDG, SQDG). To facilitate their identification, all pseudo-reactions and associated pseudo-metabolites involved in the BOFs were respectively tagged “WT526” and “MgACSBG31”. Detailed lipidomic profiles in terms of glycerolipid classes and acyl chains profiles of glycerolipids reported for “untransformed WT” and “MgACSBG#31” strains from (Billey et al., 2021) were used to calculate lipid stoichiometric coefficients. Other stoichiometric coefficients of both BOFs are based on diverse experimental and literature data. Detailed biomass composition, original data, sources and final stoichiometric coefficients are detailed in Supplemental Material and Methods and Supplemental Files S1 and S2.

### Model simulations and validation

All flux distributions were performed using open-source COBRApy package (Ebrahim et al., 2013), version 0.21.0, using cplex (version 12.8.0.0) as solver, with Python (version 3.6.13). When not otherwise stipulated, default medium was used for simulations, representing photoautotrophic growth with NO_3_ as nitrogen source with unlimited uptakes of CO_2_, O_2_, H_2_O, protons, phosphate, sulfate, magnesium, mannose, and 6-deoxy-galactose. This entailed setting the lower bounds of the corresponding exchange reactions to − 1000, as we used the usual convention of writing the exchange reactions in such way that production corresponds to positive fluxes and consumption to negative ones. Upper and lower bounds of photon and NO_3_ uptake were adjusted for each simulation. See Supplemental Material and Methods for more details.

### MEMOTE

Model quality was assessed by MEMOTE Test Suite, which calculates an independent and comparable score based on standardized tests for diverse GEM features (Lieven et al., 2020). Command version of MEMOTE (Platform Windows, Memote Version 0.12.0) was run on a conda environment with Python (version 3.6.13). MEMOTE was also run for other *Nannochloropsis* and *Microchloropsis* GEMs discussed in this study (Belcour et al., 2023; Loira et al., 2017; Shah et al., 2017). All complete MEMOTE reports are available in Supplemental Files S3-S6.

### Quantitative validation

Quantitative capability of iMgadit23 to predict growth was assessed using pFBA approach with WT BOF maximization as objective function. NO_3_ uptake rates reported by (Rafay et al., 2020); 0.119, 0.124 and 0.133 mmol .gDW^-1^.h^-1^ were used to constrain EX_no3_ reaction. Unlimited photon uptake was allowed by setting EX_photon_abs_ lower bound to −1000.

For phenotypic phase-planes analysis, couples of NO_3_-photon, and NO_3_-CO_2_ values were used to constrain NO_3_, CO_2_ and photon exchange reactions lower bound. Respectively, NO_3_, photon and CO_2_ uptake values (in mmol.gDW^-1^.h^-1^) range in [0,0.12], [0,25] and [0,1.2]. Maximum growth rate was predicted by pFBA using WT BOF as objective function.

### Qualitative validation

Qualitative growth/no-growth predictions were performed using pFBA using WT BOF as objective function. Detailed constraints applied for these simulations are available in Supplemental File S11. Model predictions were compared to in vivo growth data from literature (Fang et al., 2004; Gim et al., 2016; Kilian et al., 2011; Loira et al., 2017). In silico growth predictions were categorized as: TP, TN, FP and FN.

– TP (true positive: model simulation predicts growth while growth is observed in vivo),
– TN (true negative: model simulation predicts no growth while no growth is observed in vivo),
– FP (false positive: model simulation predicts growth while no growth is observed in vivo),
– FN (false negative: model simulation predicts no growth while growth is observed in vivo). Sensitivity, specificity, and accuracy scores were calculated according to equations (1-3).

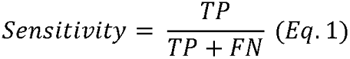

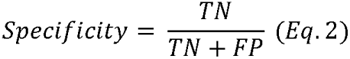

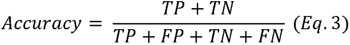

this acsbg strain-specific biomass reaction to the acsbg experimental growth rate. ACS reactions (nor any metabolic reaction) were not further constrained to model acsbg mutant.

### pFBA_WT and pFBA_ascbg flux distributions

pFBA was carried out employing the minimization of photon uptake as objective function, and constraining respectively WT BOF (BIOMASS_biomass_WT526) to 0.030 h^-1^ and acsbg mutant BOF (BIOMASS_biomass_MgACSBG31) to 0.015 h^-1^; experimental growth rates reported by (Billey et al., 2021). Unlimited NO_3_ uptake was allowed and photon uptake lower bound was constrained to −331 mmol .m^-2^.s^-1^.

### MOMA_acsbg flux distribution

MOMA was carried out employing using pFBA_WT as reference flux distribution, and constraining acsbg mutant BOF to 0.015 h^-1^. Based on experimental growth conditions, unlimited NO_3_ uptake was allowed and photon uptake lower bound was constrained to −331 mmol .m^-2^.s^-1^ (Supplemental Material and Methods). Lower and upper bounds of carbohydrates and amino acids exchange reactions were set to 0.

### Differential flux analysis

For all aforementioned flux distributions, fluxes were rounded to 10^-6^ For each iMgadit23 reaction, the change in flux between acsbg and WT was calculated as follows:

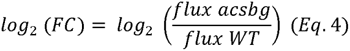

Where flux acsb9 corresponds to pFBA_acsbg or MOMA_acsbg and flux WT to pFBA_WT. log2(FC) of −0.263 and 0.263 (corresponding to a fold change of 20% in flux) were used to filter reactions with insignificant flux change. When no flux was predicted for both acsbg and WT, reaction was considered as “both_off”. When zero flux was predicted for WT and non-zero flux was predicted for acsbg, reaction was considered as activated “ON”. Finally, when non-zero flux was predicted to WT and zero flux was predicted for acsbg, reaction was considered as shut down “OFF”. Sign change in predicted fluxes was also considered by introduction of “neg_to_pos” and “pos_to_neg” labels.

### FVA_WT and FVA_acsbg

For FVA_WT, WT BOF was constrained to 0.030 h^-1^ and FVA with an optimality constraint of 100% was applied with minimization of photon uptake as objective function. Identically, for FVA_acsbg, BIOMASS_biomass_MgACSBG31 was constrained to 0.015 h^-1^ and FVA was applied with minimization of photon uptake as objective function, with an optimality constraint of 100%. For both simulations, unlimited NO_3_ uptake was allowed and photon uptake lower bound was constrained to −331 mmol .m^-2^.s^-1^. To compare the resulting minimal and maximal reaction fluxes, for each reaction i within iMgadit23, the fractional overlap (,;_i_) of the corresponding flux range between both strains was calculated as defined by (Simensen et al. 2022):

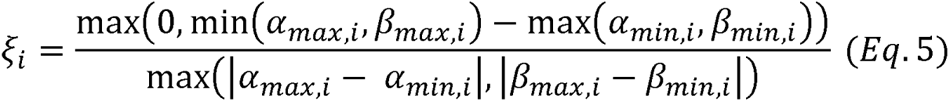

with a_min,i_ and a_max,i_ minimal and maximal fluxes for the given reaction i in FVA_WT, and {3_min,i_ and {3_max,i_ minimal and maximal fluxes in FVA_acsbg.

### iMgadit23 distribution

iMgadit23 is publicly available under a Creative Commons NonCommercial 4.0 International (CC BY-NC 4.0) license. Model and all scripts used for analyses presented in this study are available in a GitHub repository (https://github.com/Total-RD/Mgaditana-GEM), following the standard-GEM format (https://github.com/MetabolicAtlas/standard-GEM; (Anton et al., 2023). Model reconstruction is provided in SBML, JSON, MAT and XLSX formats compatible with the metabolic engineering platforms and computational tools for metabolic model simulations (COBRA, Escher and others) (Supplemental File S12 and https://github.com/Total-RD/Mgaditana-GEM). Detailed 2D-pathway map of iMgadit23 content is available as JSON file compatible with Escher (King et al., 2015) (https://github.com/Total-RD/Mgaditana-GEM).

## Supporting information

Supplemental Results

## Supporting information

### Supplemental Results: Figures

Figure S1. Photosynthesis modeling and impact of PSQUANTUM in iMgadit23.

Figure S2. Results of pFBA differential flux analysis for lipid metabolism.

Figure S3. Results of MOMA differential flux analysis for lipid metabolism.

### Tables

Table S1. log2(pFBA_acsbg/pFBA_WT) results. Table S2. log2(MOMA_acsbg/pFBA_WT) results.

## Supplemental Material and Methods

### Figures

Figure S4. Specific growth rate and quantum yield used for iMgadit23 validation.

Figure S5. Linear correlation between photon uptake and optimal growth rate predicted by pFBA.

### Tables

Table S3. Fatty acid nomenclature. Table S4. Glycerolipid class acronyms.

Table S5. TAGL coding genes in NagaB31_1.0. Table S6. Default constraints within iMgadit23.

Table S7. Nitrate uptake data used to constrain iMgadit23 for quantitative validation.

Supplemental files: Supp_File_S1_BOF_WT526_c.xlsx

Supp_File_S2_BOF_MgACSBG31_c.xlsx

Supp_File_S3_RES_MEMOTE_iMgadit23.html

Supp_File_S4_RES_Memote_AucomeNannochloropsisGaditana.html

Supp_File_S5_RES_Memote_iNS937.html

Supp_file_S6_RES_Memote_iRJ1321_auto.html

Supp_File_S7_Test_growth_KO_ACS_rxn.xlsx

Supp_File_S8_ACSBG_compare_pFBA_MOMA_supp_to_scatter.xlsx

Supp_File_S9_Bilan_ACSBG_fluxes.xlsx

Supp_File_S10_ACSBG_results_TAG_biosynthesis.xlsx

Supp_File_S11_Media_composition_qualitative_validation.csv Supp_File_S12_iMgadit23.xlsx

## Acknowledgments and Funding

The Authors thank Abril Lansoy from TotalEnergies and Florian Delrue (CEA Cadarache) for their supports, the SilicoLife team for their manual validation of the functional annotations and for their collaboration in the rest of the model reconstruction. This work was supported by the ANRT via a Cifre PhD (TotalEnergies&CEA) n°2020/1063 and by GRAL, the Grenoble Alliance for Integrated Structural & Cell Biology, a program from the Chemistry Biology Health Graduate School of University Grenoble Alpes (ANR-17-EURE-0003).

## Author Contributions

CDT, SC, BP, RC, PV performed research; GF, EM, EB, GC, MD and JJ designed the research and analyzed data; CDT and JJ wrote the paper, all the authors reviewed the paper.

## Notes

https://github.com/Total-RD/Mgaditana-GEM

### Competing Interest Statement

The authors have declared no competing interest.

### Summary of Updates

Introduction and Material and Method updated to clarify

## References

Alboresi, A., Perin, G., Vitulo, N., Diretto, G., Block, M., Jouhet, J., Meneghesso, A., Valle, G., Giuliano, G., Maréchal, E., Morosinotto, T., 2016. Light Remodels Lipid Biosynthesis in *Nannochloropsis gaditana* by Modulating Carbon Partitioning between Organelles. Plant Physiol 171, 2468–2482. 10.1104/pp.16.00599

Al-Hoqani, U., Young, R., Purton, S., 2017. The biotechnological potential of Nannochloropsis. Perspectives in Phycology 1–15. 10.1127/pip/2016/0065

Allen, J.F., 2002. Photosynthesis of ATP—Electrons, Proton Pumps, Rotors, and Poise. Cell 110, 273–276. 10.1016/S0092-8674(02)00870-X

Altschul, S.F., Gish, W., Miller, W., Myers, E.W., Lipman, D.J., 1990. Basic local alignment search tool. Journal of Molecular Biology 215, 403–410. 10.1016/S0022-2836(05)80360-2

Anton, M., Almaas, E., Benfeitas, R., Benito-Vaquerizo, S., Blank, L.M., Dräger, A., Hancock, J.M., Kittikunapong, C., König, M., Li, F., Liebal, U.W., Lu, H., Ma, H., Mahadevan, R., Mardinoglu, A., Nielsen, J., Nogales, J., Pagni, M., Papin, J.A., Patil, K.R., Price, N.D., Robinson, J.L., Sánchez, B.J., Suarez-Diez, M., Sulheim, S., Svensson, L.T., Teusink, B., Vongsangnak, W., Wang, H., Zeidan, A.A., Kerkhoven, E.J., 2023. standard-GEM: standardization of open-source genome-scale metabolic models. 10.1101/2023.03.21.512712

Belcour, A., Got, J., Aite, M., Delage, L., Collén, J., Frioux, C., Leblanc, C., Dittami, S.M., Blanquart, S., Markov, G.V., Siegel, A., 2023. Inferring and comparing metabolism across heterogeneous sets of annotated genomes using AuCoMe. Genome Res 33, 972–987. 10.1101/gr.277056.122

Billey, E., Magneschi, L., Leterme, S., Bedhomme, M., Andres-Robin, A., Poulet, L., Michaud, M., Finazzi, G., Dumas, R., Crouzy, S., Laueffer, F., Fourage, L., Rébeillé, F., Amato, A., Collin, S., Jouhet, J., Maréchal, E., 2021. Characterization of the Bubblegum acyl-CoA synthetase of *Microchloropsis gaditana*. Plant Physiology 185, 815–835. 10.1093/plphys/kiaa110

Bo, D.D., Magneschi, L., Bedhomme, M., Billey, E., Deragon, E., Storti, M., Menneteau, M., Richard, C., Rak, C., Lapeyre, M., Lembrouk, M., Conte, M., Gros, V., Tourcier, G., Giustini, C., Falconet, D., Curien, G., Allorent, G., Petroutsos, D., Laeuffer, F., Fourage, L., Jouhet, J., Maréchal, E., Finazzi, G., Collin, S., 2021. Consequences of Mixotrophy on Cell Energetic Metabolism in *Microchloropsis gaditana* Revealed by Genetic Engineering and Metabolic Approaches. Frontiers in Plant Science 12.

Boyle, N.R., Morgan, J.A., 2009. Flux balance analysis of primary metabolism in Chlamydomonas reinhardtii. BMC Syst Biol 3, 4. 10.1186/1752-0509-3-4

Chu, W.-L., 2012. Biotechnological applications of microalgae. IeJSME 6, S24–S37. 10.56026/imu.6.Suppl1.S24

Correa, S.M., Fernie, A.R., Nikoloski, Z., Brotman, Y., 2020. Towards model-driven characterization and manipulation of plant lipid metabolism. Progress in Lipid Research 80, 101051. 10.1016/j.plipres.2020.101051

Corteggiani Carpinelli, E., Telatin, A., Vitulo, N., Forcato, C., D’Angelo, M., Schiavon, R., Vezzi, A., Giacometti, G.M., Morosinotto, T., Valle, G., 2014. Chromosome scale genome assembly and transcriptome profiling of *Nannochloropsis gaditana* in nitrogen depletion. Mol Plant 7, 323–335. 10.1093/mp/sst120

Curien, G., Flori, S., Villanova, V., Magneschi, L., Giustini, C., Forti, G., Matringe, M., Petroutsos, D., Kuntz, M., Finazzi, G., 2016. The Water to Water Cycles in Microalgae. Plant and Cell Physiology 57, 1354–1363. 10.1093/pcp/pcw048

Dolch, L.-J., Rak, C., Perin, G., Tourcier, G., Broughton, R., Leterrier, M., Morosinotto, T., Tellier, F., Faure, J.-D., Falconet, D., Jouhet, J., Sayanova, O., Beaudoin, F., Maréchal, E., 2017. A Palmitic Acid Elongase Affects Eicosapentaenoic Acid and Plastidial Monogalactosyldiacylglycerol Levels in Nannochloropsis. Plant Physiol 173, 742–759. 10.1104/pp.16.01420

Ebrahim, A., Lerman, J.A., Palsson, B.O., Hyduke, D.R., 2013. COBRApy: COnstraints-Based Reconstruction and Analysis for Python. BMC Syst Biol 7, 74. 10.1186/1752-0509-7-74

Eddy, S.R., 1998. Profile hidden Markov models. Bioinformatics 14, 755–763. 10.1093/bioinformatics/14.9.755

Fang, X., Wei, C., Zhao-Ling, C., Fan, O., 2004. Effects of organic carbon sources on cell growth and eicosapentaenoic acid content of Nannochloropsis sp. J Appl Phycol 16, 499–503. 10.1007/s10811-004-5520-1

Fawley, M.W., Jameson, I., Fawley, K.P., 2015. The phylogeny of the genus Nannochloropsis (Monodopsidaceae, Eustigmatophyceae), with descriptions of N. australis sp. nov. and Microchloropsis gen. nov. Phycologia 54, 545–552. 10.2216/15-60.1

Fu, W., Nelson, D.R., Mystikou, A., Daakour, S., Salehi-Ashtiani, K., 2019. Advances in microalgal research and engineering development. Current Opinion in Biotechnology, Tissue, Cell and Pathway Engineering 59, 157–164. 10.1016/j.copbio.2019.05.013

Gentile, M.P., Blanch, H.W., 2001. Physiology and xanthophyll cycle activity of *Nannochloropsis gaditana*. Biotechnol Bioeng 75, 1–12. 10.1002/bit.1158

Gim, G.H., Ryu, J., Kim, M.J., Kim, P.I., Kim, S.W., 2016. Effects of carbon source and light intensity on the growth and total lipid production of three microalgae under different culture conditions. J Ind Microbiol Biotechnol 43, 605–616. 10.1007/s10295-016-1741-y

Gu, C., Kim, G.B., Kim, W.J., Kim, H.U., Lee, S.Y., 2019. Current status and applications of genome-scale metabolic models. Genome Biol 20, 121. 10.1186/s13059-019-1730-3

Gudmundsson, S., Thiele, I., 2010. Computationally efficient flux variability analysis. BMC Bioinformatics 11, 489. 10.1186/1471-2105-11-489

Hahn, A., Vonck, J., Mills, D.J., Meier, T., Kühlbrandt, W., 2018. Structure, mechanism, and regulation of the chloroplast ATP synthase. Science 360, eaat4318. 10.1126/science.aat4318

Heirendt, L., Arreckx, S., Pfau, T., Mendoza, S.N., Richelle, A., Heinken, A., Haraldsdóttir, H.S., Wachowiak, J., Keating, S.M., Vlasov, V., Magnusdóttir, S., Ng, C.Y., Preciat, G., Žagare, A., Chan, S.H.J., Aurich, M.K., Clancy, C.M., Modamio, J., Sauls, J.T., Noronha, A., Bordbar, A., Cousins, B., El Assal, D.C., Valcarcel, L.V., Apaolaza, I., Ghaderi, S., Ahookhosh, M., Ben Guebila, M., Kostromins, A., Sompairac, N., Le, H.M., Ma, D., Sun, Y., Wang, L., Yurkovich, J.T., Oliveira, M.A.P., Vuong, P.T., El Assal, L.P., Kuperstein, I., Zinovyev, A., Hinton, H.S., Bryant, W.A., Aragón Artacho, F.J., Planes, F.J., Stalidzans, E., Maass, A., Vempala, S., Hucka, M., Saunders, M.A., Maranas, C.D., Lewis, N.E., Sauter, T., Palsson, B.Ø., Thiele, I., Fleming, R.M.T., 2019. Creation and analysis of biochemical constraint-based models using the COBRA Toolbox v.3.0. Nat Protoc 14, 639–702. 10.1038/s41596-018-0098-2

Hoffmann, D.Y., Shachar-Hill, Y., 2023. Do betaine lipids replace phosphatidylcholine as fatty acid editing hubs in microalgae? Front Plant Sci 14, 1077347. 10.3389/fpls.2023.1077347

Jouhet, J., Lupette, J., Clerc, O., Magneschi, L., Bedhomme, M., Collin, S., Roy, S., Maréchal, E., Rébeillé, F., 2017. LC-MS/MS versus TLC plus GC methods: Consistency of glycerolipid and fatty acid profiles in microalgae and higher plant cells and effect of a nitrogen starvation. PLOS ONE 12, e0182423. 10.1371/journal.pone.0182423

Kanehisa, M., Goto, S., 2000. KEGG: Kyoto Encyclopedia of Genes and Genomes. Nucleic Acids Research 28, 27–30. 10.1093/nar/28.1.27

Kareya, M.S., Mariam, I., Shaikh, K.M., Nesamma, A.A., Jutur, P.P., 2020. Photosynthetic Carbon Partitioning and Metabolic Regulation in Response to Very-Low and High CO2 in *Microchloropsis gaditana* NIES 2587. Front Plant Sci 11, 981. 10.3389/fpls.2020.00981

Keeling, P.J., 2004. Diversity and evolutionary history of plastids and their hosts. Am J Bot 91, 1481–1493. 10.3732/ajb.91.10.1481

Kilian, O., Benemann, C.S.E., Niyogi, K.K., Vick, B., 2011. High-efficiency homologous recombination in the oil-producing alga Nannochloropsis sp. Proc Natl Acad Sci U S A 108, 21265–21269. 10.1073/pnas.1105861108

Kim, C.W., Sung, M.-G., Nam, K., Moon, M., Kwon, J.-H., Yang, J.-W., 2014. Effect of monochromatic illumination on lipid accumulation of *Nannochloropsis gaditana* under continuous cultivation. Bioresour Technol 159, 30–35. 10.1016/j.biortech.2014.02.024

King, Z.A., Dräger, A., Ebrahim, A., Sonnenschein, N., Lewis, N.E., Palsson, B.O., 2015. Escher: A Web Application for Building, Sharing, and Embedding Data-Rich Visualizations of Biological Pathways. PLoS Comput Biol 11, e1004321. 10.1371/journal.pcbi.1004321

Kong, F., Romero, I.T., Warakanont, J., Li-Beisson, Y., 2018. Lipid catabolism in microalgae. New Phytol 218, 1340–1348. 10.1111/nph.15047

Levering, J., Broddrick, J., Dupont, C.L., Peers, G., Beeri, K., Mayers, J., Gallina, A.A., Allen, A.E., Palsson, B.O., Zengler, K., 2016. Genome-Scale Model Reveals Metabolic Basis of Biomass Partitioning in a Model Diatom. PLOS ONE 11, e0155038. 10.1371/journal.pone.0155038

Lewis, N.E., Hixson, K.K., Conrad, T.M., Lerman, J.A., Charusanti, P., Polpitiya, A.D., Adkins, J.N., Schramm, G., Purvine, S.O., Lopez-Ferrer, D., Weitz, K.K., Eils, R., König, R., Smith, R.D., Palsson, B.Ø., 2010. Omic data from evolved E. coli are consistent with computed optimal growth from genome-scale models. Mol Syst Biol 6, 390. 10.1038/msb.2010.47

Li-Beisson, Y., Thelen, J.J., Fedosejevs, E., Harwood, J.L., 2019. The lipid biochemistry of eukaryotic algae. Progress in Lipid Research 74, 31–68. 10.1016/j.plipres.2019.01.003

Lieven, C., Beber, M.E., Olivier, B.G., Bergmann, F.T., Ataman, M., Babaei, P., Bartell, J.A., Blank, L.M., Chauhan, S., Correia, K., Diener, C., Dräger, A., Ebert, B.E., Edirisinghe, J.N., Faria, J.P., Feist, A.M., Fengos, G., Fleming, R.M.T., García-Jiménez, B., Hatzimanikatis, V., van Helvoirt, W., Henry, C.S., Hermjakob, H., Herrgård, M.J., Kaafarani, A., Kim, H.U., King, Z., Klamt, S., Klipp, E., Koehorst, J.J., König, M., Lakshmanan, M., Lee, D.-Y., Lee, S.Y., Lee, S., Lewis, N.E., Liu, F., Ma, H., Machado, D., Mahadevan, R., Maia, P., Mardinoglu, A., Medlock, G.L., Monk, J.M., Nielsen, J., Nielsen, L.K., Nogales, J., Nookaew, I., Palsson, B.O., Papin, J.A., Patil, K.R., Poolman, M., Price, N.D., Resendis-Antonio, O., Richelle, A., Rocha, I., Sánchez, B.J., Schaap, P.J., Malik Sheriff, R.S., Shoaie, S., Sonnenschein, N., Teusink, B., Vilaça, P., Vik, J.O., Wodke, J.A.H., Xavier, J.C., Yuan, Q., Zakhartsev, M., Zhang, C., 2020. MEMOTE for standardized genome-scale metabolic model testing. Nat Biotechnol 38, 272–276. 10.1038/s41587-020-0446-y

Lima, S., Brucato, A., Caputo, G., Schembri, L., Scargiali, F., 2022. Modelling *Nannochloropsis gaditana* Growth in Reactors with Different Geometries, Determination of Kinetic Parameters and Biochemical Analysis in Response to Light Intensity. Applied Sciences 12, 5776. 10.3390/app12125776

Loira, N., Mendoza, S., Paz Cortés, M., Rojas, N., Travisany, D., Genova, A.D., Gajardo, N., Ehrenfeld, N., Maass, A., 2017. Reconstruction of the microalga Nannochloropsis salina genome-scale metabolic model with applications to lipid production. BMC Syst Biol 11, 66. 10.1186/s12918-017-0441-1

Ma, X., Mi, Y., Zhao, C., Wei, Q., 2022. A comprehensive review on carbon source effect of microalgae lipid accumulation for biofuel production. Sci Total Environ 806, 151387. 10.1016/j.scitotenv.2021.151387

Ma, X.-N., Chen, T.-P., Yang, B., Liu, J., Chen, F., 2016. Lipid Production from Nannochloropsis. Mar Drugs 14, 61. 10.3390/md14040061

Min, K.-T., Benzer, S., 1999. Preventing Neurodegeneration in the Drosophila Mutant bubblegum. Science 284, 1985–1988. 10.1126/science.284.5422.1985

Mühlroth, A., Winge, P., El Assimi, A., Jouhet, J., Maréchal, E., Hohmann-Marriott, M.F., Vadstein, O., Bones, A.M., 2017. Mechanisms of Phosphorus Acquisition and Lipid Class Remodeling under P Limitation in a Marine Microalga. Plant Physiology 175, 1543–1559. 10.1104/pp.17.00621

Muñoz, C.F., Südfeld, C., Naduthodi, M.I.S., Weusthuis, R.A., Barbosa, M.J., Wijffels, R.H., D’Adamo, S., 2021. Genetic engineering of microalgae for enhanced lipid production. Biotechnol Adv 52, 107836. 10.1016/j.biotechadv.2021.107836

Norsigian, C.J., Pusarla, N., McConn, J.L., Yurkovich, J.T., Dräger, A., Palsson, B.O., King, Z., 2020. BiGG Models 2020: multi-strain genome-scale models and expansion across the phylogenetic tree. Nucleic Acids Res 48, D402–D406. 10.1093/nar/gkz1054

Orth, J.D., Thiele, I., Palsson, B.Ø., 2010. What is flux balance analysis? Nat Biotechnol 28, 245–248. 10.1038/nbt.1614

Pereira, B., Miguel, J., Vilaça, P., Soares, S., Rocha, I., Carneiro, S., 2018. Reconstruction of a genome-scale metabolic model for Actinobacillus succinogenes 130Z. BMC Systems Biology 12, 61. 10.1186/s12918-018-0585-7

Petroutsos, D., Amiar, S., Abida, H., Dolch, L.-J., Bastien, O., Rébeillé, F., Jouhet, J., Falconet, D., Block, M.A., McFadden, G.I., Bowler, C., Botté, C., Maréchal, E., 2014. Evolution of galactoglycerolipid biosynthetic pathways--from cyanobacteria to primary plastids and from primary to secondary plastids. Prog Lipid Res 54, 68–85. 10.1016/j.plipres.2014.02.001

Pham, N., 2016. Genome-scale constraint-based metabolic modeling and analysis of Nannochloropsis Sp. (Master thesis). NTNU.

Poliner, E., Panchy, N., Newton, L., Wu, G., Lapinsky, A., Bullard, B., Zienkiewicz, A., Benning, C., Shiu, S.-H., Farré, E.M., 2015. Transcriptional coordination of physiological responses in Nannochloropsis oceanica CCMP1779 under light/dark cycles. Plant J 83, 1097–1113. 10.1111/tpj.12944

Rafay, R., Uratani, J.M., Hernandez, H.H., Rodríguez, J., 2020. Growth and Nitrate Uptake in *Nannochloropsis gaditana* and Tetraselmis chuii Cultures Grown in Sequential Batch Reactors. Frontiers in Marine Science 7.

Ren, M., Ogden, K., 2014. Cultivation of *Nannochloropsis gaditana* on mixtures of nitrogen sources. Environmental Progress & Sustainable Energy 33, 551–555. 10.1002/ep.11818

Rizwan, M., Mujtaba, G., Memon, S.A., Lee, K., Rashid, N., 2018. Exploring the potential of microalgae for new biotechnology applications and beyond: A review. Renewable and Sustainable Energy Reviews 92, 394–404. 10.1016/j.rser.2018.04.034

Rock, A., Novoveska, L., Green, D., 2021. Synthetic biology is essential to unlock commercial biofuel production through hyper lipid-producing microalgae: a review. Applied Phycology 2, 2021. 10.1080/26388081.2021.1886872

Rowe, E., Palsson, B.O., King, Z.A., 2018. Escher-FBA: a web application for interactive flux balance analysis. BMC Systems Biology 12, 84. 10.1186/s12918-018-0607-5

Saier, M.H., Jr, Reddy, V.S., Tamang, D.G., Västermark, Å., 2014. The Transporter Classification Database. Nucleic Acids Research 42, D251–D258. 10.1093/nar/gkt1097

Sajjadi, B., Chen, W.-Y., Raman Abdul.Aziz. A., Ibrahim, S., 2018. Microalgae lipid and biomass for biofuel production: A comprehensive review on lipid enhancement strategies and their effects on fatty acid composition. Renewable and Sustainable Energy Reviews 97, 200–232. 10.1016/j.rser.2018.07.050

Sánchez, B.J., Li, F., Kerkhoven, E.J., Nielsen, J., 2019. SLIMEr: probing flexibility of lipid metabolism in yeast with an improved constraint-based modeling framework. BMC Systems Biology 13, 4. 10.1186/s12918-018-0673-8

Segrè, D., Vitkup, D., Church, G.M., 2002. Analysis of optimality in natural and perturbed metabolic networks. Proc Natl Acad Sci U S A 99, 15112–15117. 10.1073/pnas.232349399

Shah, A.R., Ahmad, A., Srivastava, S., Jaffar Ali, B.M., 2017. Reconstruction and analysis of a genome-scale metabolic model of *Nannochloropsis gaditana*. Algal Research 26, 354–364. 10.1016/j.algal.2017.08.014

Simionato, D., Block, M.A., La Rocca, N., Jouhet, J., Maréchal, E., Finazzi, G., Morosinotto, T., 2013. The Response of *Nannochloropsis gaditana* to Nitrogen Starvation Includes De Novo Biosynthesis of Triacylglycerols, a Decrease of Chloroplast Galactolipids, and Reorganization of the Photosynthetic Apparatus. Eukaryotic Cell 12, 665–676. 10.1128/ec.00363-12

Song, S.-Y., Kato, C., Adachi, E., Moriya-Sato, A., Inagawa-Ogashiwa, M., Umeda, R., Hashimoto, N., 2007. Expression of an acyl-CoA synthetase, lipidosin, in astrocytes of the murine brain and its up-regulation during remyelination following cuprizone-induced demyelination. J Neurosci Res 85, 3586–3597. 10.1002/jnr.21456

Spolaore, P., Joannis-Cassan, C., Duran, E., Isambert, A., 2006. Commercial applications of microalgae. J Biosci Bioeng 101, 87–96. 10.1263/jbb.101.87

Steinberg, S.J., Morgenthaler, J., Heinzer, A.K., Smith, K.D., Watkins, P.A., 2000. Very Long-chain Acyl-CoA Synthetases: HUMAN “BUBBLEGUM” REPRESENTS A NEW FAMILY OF PROTEINS CAPABLE OF ACTIVATING VERY LONG-CHAIN FATTY ACIDS*. Journal of Biological Chemistry 275, 35162– 35169. 10.1074/jbc.M006403200

Tibocha-Bonilla, J.D., Zuñiga, C., Godoy-Silva, R.D., Zengler, K., 2018. Advances in metabolic modeling of oleaginous microalgae. Biotechnol Biofuels 11, 241. 10.1186/s13068-018-1244-3

UniProt Consortium, T., 2018. UniProt: the universal protein knowledgebase. Nucleic Acids Research 46, 2699. 10.1093/nar/gky092

Vieler, A., Wu, G., Tsai, C.-H., Bullard, B., Cornish, A.J., Harvey, C., Reca, I.-B., Thornburg, C., Achawanantakun, R., Buehl, C.J., Campbell, M.S., Cavalier, D., Childs, K.L., Clark, T.J., Deshpande, R., Erickson, E., Armenia Ferguson, A., Handee, W., Kong, Q., Li, X., Liu, B., Lundback, S., Peng, C., Roston, R.L., Sanjaya, null, Simpson, J.P., Terbush, A., Warakanont, J., Zäuner, S., Farre, E.M., Hegg, E.L., Jiang, N., Kuo, M.-H., Lu, Y., Niyogi, K.K., Ohlrogge, J., Osteryoung, K.W., Shachar-Hill, Y., Sears, B.B., Sun, Y., Takahashi, H., Yandell, M., Shiu, S.-H., Benning, C., 2012. Genome, functional gene annotation, and nuclear transformation of the heterokont oleaginous alga Nannochloropsis oceanica CCMP1779. PLoS Genet 8, e1003064. 10.1371/journal.pgen.1003064

Witting, M., 2020. Suggestions for Standardized Identifiers for Fatty Acyl Compounds in Genome Scale Metabolic Models and Their Application to the WormJam Caenorhabditis elegans Model. Metabolites 10, 130. 10.3390/metabo10040130

Ye, C., Wei, X., Shi, T., Sun, X., Xu, N., Gao, C., Zou, W., 2022. Genome-scale metabolic network models: from first-generation to next-generation. Appl Microbiol Biotechnol 106, 4907–4920. 10.1007/s00253-022-12066-y

Zhang, D., Yan, F., Sun, Z., Zhang, Q., Xue, S., Cong, W., 2014. On-line modeling intracellular carbon and energy metabolism of Nannochloropsis sp. in nitrogen-repletion and nitrogen-limitation cultures. Bioresour Technol 164, 86–92. 10.1016/j.biortech.2014.04.083

