## Supplemental Material and Methods for "A new genome-scale metabolic model of oleaginous microalgae with refined lipid metabolism elucidates *Microchloropsis gaditana* mutant phenotypes"

Lipid nomenclature in iMgadit23

In order to standardize the nomenclature within iMgadit23 and make it clearer and more understandable, the identifiers for lipid metabolites have been standardized.

**Fatty acid nomenclature**

The nomenclature used is based on the abbreviated chemical nomenclature (C: D number of carbons : number of double bonds) adapted to the BiGG identifier specifications.

ID = fatty acid class + acyl chain C:D (without “:”)

Examples of different classes of fatty acids and fatty acid derivatives are shown in theTable S3.

**Table S3. Fatty acid nomenclature.**

| **Fatty acid class** | **Abreviation** | **Suffix** | **Exemple (16:0)** |
| --- | --- | --- | --- |
| Fatty acid | c | / | **c**160 |
| Fatty acid -ACP (**A**cyl **C**arrier **P**rotein) | c | ACP | **c**160_**ACP** |
| Acyl-CoA | c | coa | **c**160_**coa** |
| Fatty acid-FAS1 | c | FAS1 | **c**160_**FAS1** |
| 3-oxoacyl-CoA | 3oc | coa | **3oc**160_**coa** |
| 3-hydroxyacyl-CoA | 3hc | coa | **3hc**160_**coa** |
| Trans-2-enoyl-CoA | c | 2e_coa | **c**161_**2e**_**coa** |

**Glycerolipid nomenclature**

Several nomenclatures exist to designate glycerolipids. These nomenclatures specify the class of glycerolipid and the number of carbons and double bonds it contains, depending on the level of detail available. When the fatty acid composition is known, this can be detailed by listing the position of the fatty acids on the glycerol backbone in the order*"(sn-1/sn-2/sn-3)"*.

ID = "glycerolipid class" + "fatty acid *sn-1*" + "_" + " fatty acid *sn-2*"

TAGs harboring three acyl chains are identified :

ID = tag + "fatty acid *sn-1*" + "_" + " fatty acid *sn-2*" + "_" + " fatty acid *sn-3*"

The acronyms used for the different classes of glycerolipids are shown in Table S4.

**Table S4. Glycerolipid class acronyms.**

| **Glycerolipid Class** | **Acronym** |
| --- | --- |
| Phosphatidylcholine | pc |
| Phosphatidylethanolamine | pe |
| Phosphatidylinositol | pi |
| Phosphatidylglycerol | pg |
| Monogalactosyldiacylglycerol | mgdg |
| Digalactosyldiacylglycerol | dgdg |
| Sulfoquinovosyldiacylglycerol | sqdg |
| Diacylglycerol-N,N,N-trimethylhomoserine | dgts |
| Monoacylglycerol | mag |
| Diacylglycerol | dag |
| Triacylglycerol | tag |
| Lysophosphatidic acid | sn1_glyc3p |
| *sn-1* lyso-phosphatidylcholine | sn1_pc |
| *sn-1* lyso-diacylglycerol-N,N,N-trimethylhomoserine | sn1_dgts |
| *sn-2* lyso-monogalactosyldiacylglycerol | sn2_mgdg |

Addition of lipid degradation and acyl editing in iMgadit23

Acyl editing and glycerolipid degradation pathways addition in iMgadit23 was based on the following arguments.

**TAG/DAG/MAG degradation**

Since several genes of M. gaditana B-31 genome are annotated as TAG or MAG lipases, TAG, DAG and MAG degradation was added to iMgadit23.

**TAG lipases:**

Several putative TAG lipases (Naga_100677g1, Naga_100055g28, Naga_102685g1, Naga_100052g28, Naga_100718g2, Naga_100045g8, Naga_100016g73, Naga_100039g12, Naga_100171g1, Naga_100104g14, Naga_100529g6, Naga_100043g24, Naga_100101g8, Naga_100133g7, Naga_100426g5, Naga_100091g1, Naga_100271g1, Naga_100008g63, Naga_100101g12, Naga_100011g88, Naga_100013g62, Naga_100530g1, Naga_100144g12, Naga_100241g4, Naga_100247g4, Naga_100046g19, Naga_100012g35) were identified in *M. gaditana* genome (Table S4).

**Table S5. TAGL coding genes in NagaB31_1.0.**

| Gene | Identified as homologous to a TAGL | Associated to TAG degradation in Shah GSM model | Annotated as "Lipase, class 3" in NagaB31_1.0 | Annotated as "triacylglycerol lipase" in NagaB31_1.0 | Annotated as lipase in NagaB31_1.0 |
| --- | --- | --- | --- | --- | --- |
| Naga_100677g1 | homologous to Arabidopsis SDP1 and Phaeodactylum Tgl1 | √ |  |  |  |
| Naga_100055g28 | homologous to Arabidopsis SDP1 and Phaeodactylum Tgl1 | √ |  |  |  |
| Naga_102685g1 | TAGL-Like-A |  |  |  |  |
| Naga_100052g28 | Homologous to Human PNPLA3 |  |  |  |  |
| Naga_100247g4 |  | √ |  |  | PNPLA domain |
| Naga_100171g1 |  | √ | √ |  |  |
| Naga_100104g14 |  | √ | √ |  |  |
| Naga_100529g6 |  | √ | √ |  |  |
| Naga_100043g24 |  | √ | √ |  |  |
| Naga_100101g8 |  | √ | √ |  |  |
| Naga_100133g7 |  | √ | √ |  |  |
| Naga_100426g5 |  | √ | √ |  |  |
| Naga_100091g1 |  | √ | √ |  |  |
| Naga_100271g1 |  | √ | √ |  |  |
| Naga_100008g63 |  | √ | √ |  |  |
| Naga_100101g12 |  | √ | √ |  |  |
| Naga_100144g12 |  | √ | √ |  |  |
| Naga_100045g8 |  | √ |  | √ |  |
| Naga_100013g62 |  | √ |  | √ |  |
| Naga_100011g88 |  | √ |  |  | √ |
| Naga_100530g1 |  | √ |  |  | √ |
| Naga_100241g4 |  | √ |  |  | √ |
| Naga_100046g19 |  | √ |  |  | √ |
| Naga_100012g35 |  | √ |  |  | √ |
| Naga_100718g2 |  | √ |  |  | √ |
| Naga_100016g73 |  | √ |  |  | √ |
| Naga_100039g12 |  | √ |  |  | √ |

**MAG lipases:**

Putative monoacylglycerol (MAG) lipases (Naga_100049g29 and Naga_100130g4) were identified in *M. gaditana* genome.

**Plastid Galactoglycerolipid Degradation / Acyl editing on MGDG**

Complete acyl editing cycle of MGDG was reported for *Chlamydomonas reinhardtii.* Plastid Galactoglycerolipid Degradation 1 (PGD1) catalyzes the hydrolysis of acyl chains at the *sn-1* position of the glycerol backbone of the plastidial MGDG forming a lyso-MGDG (Li, Benning and Kuo, 2012). Recently, CrLAT1 enzyme catalyzing the reacylation of lyso-MGDG was characterized in *C. reinhardtii* (Hoffmann and Shachar-Hill, 2023). Lyso-MGDG were also identified in *Emiliana huxleyi* (Malitsky *et al.*, 2016).

**Phospholipases**

One gene of in *M. gaditana* genome: Naga_100454g3 is annotated as phospholipase D and contains a PLD domain. This gene was associated with hydrolysis of PC, producing PA and a free choline group.

**Acyl editing on PC and DGTS**

Evidence of acyl editing on PC or DGTS was reported for microalgae in general (Hoffmann and Shachar-Hill, 2023).

**Obtention of (Raso et al. 2012) data for photosynthesis validation**

WebPlotDigitizer (<https://apps.automeris.io/wpd/>) was used to extract specific growth rate and quantum yields from (Raso et al. 2012) data (Figure S4)

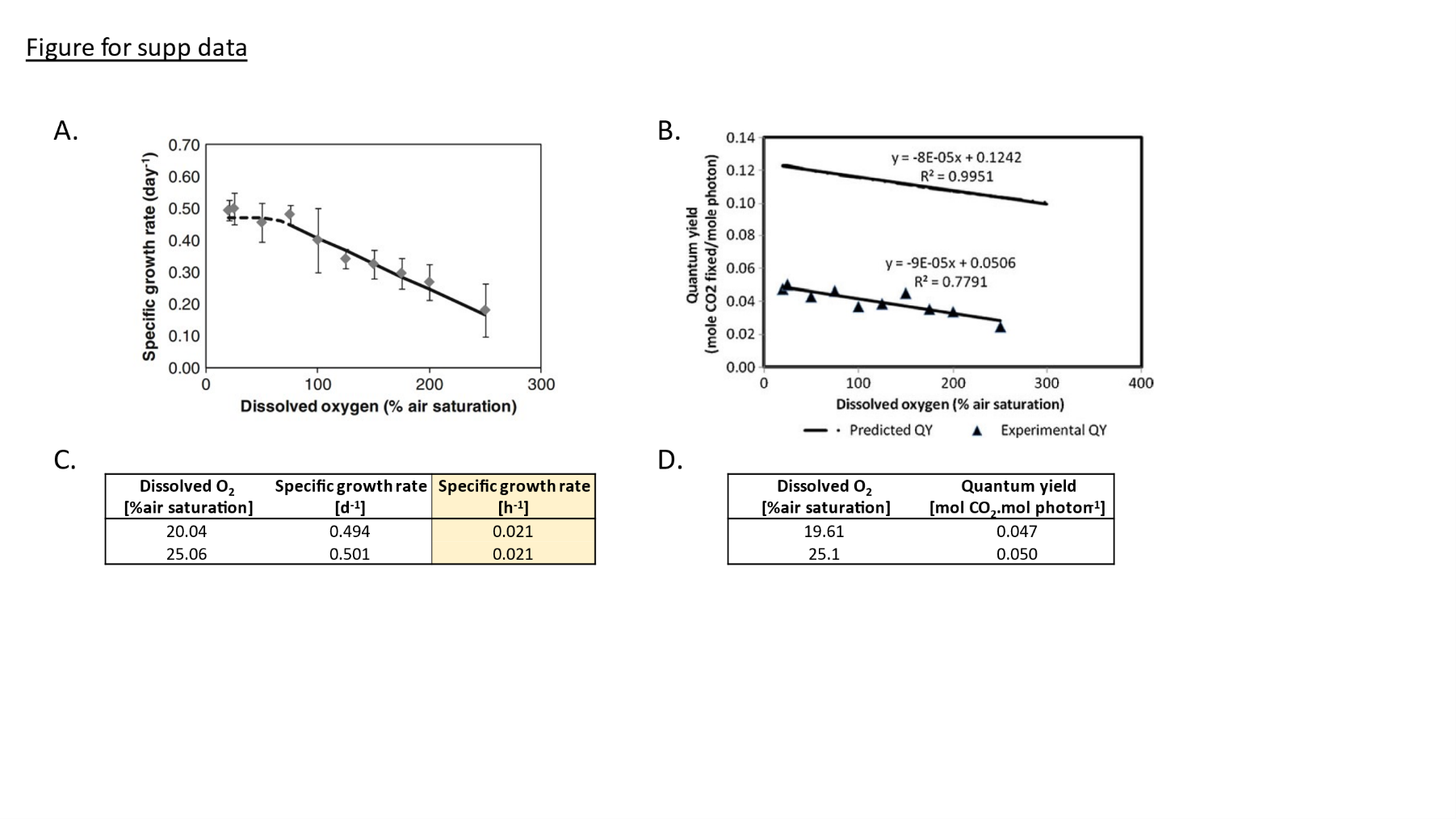

**Figure S4. Specific growth rate and quantum yield used for iMgadit23 validation.**

*(A) Effects of oxygen concentration on the specific growth rate (day^−1^) of Nannochloropsis sp. at an irradiance of 100 μmol_photons_.m^−2^.s^−1^* ***from (Raso et al. 2012); Figure 3)*** *(B) Effect of oxygen concentration on the in vivo quantum yield of Nannochloropsis sp. measured at an irradiance of 100 μmol_photons_.m^−2^.s^−1^* ***from (Raso et al. 2012); Figure 6)****. Numerical specific growth rates (C) and quantum yields (D) for 20% and 25% dissolved O_2_ were extracted with WebPlotDigitizer.*

In default medium conditions, photons are the only limiting "substrate". Consequently, maximal growth rate predicted with pFBA is linearly correlated to maximal photon uptake (r^2^=1; Figure S5).

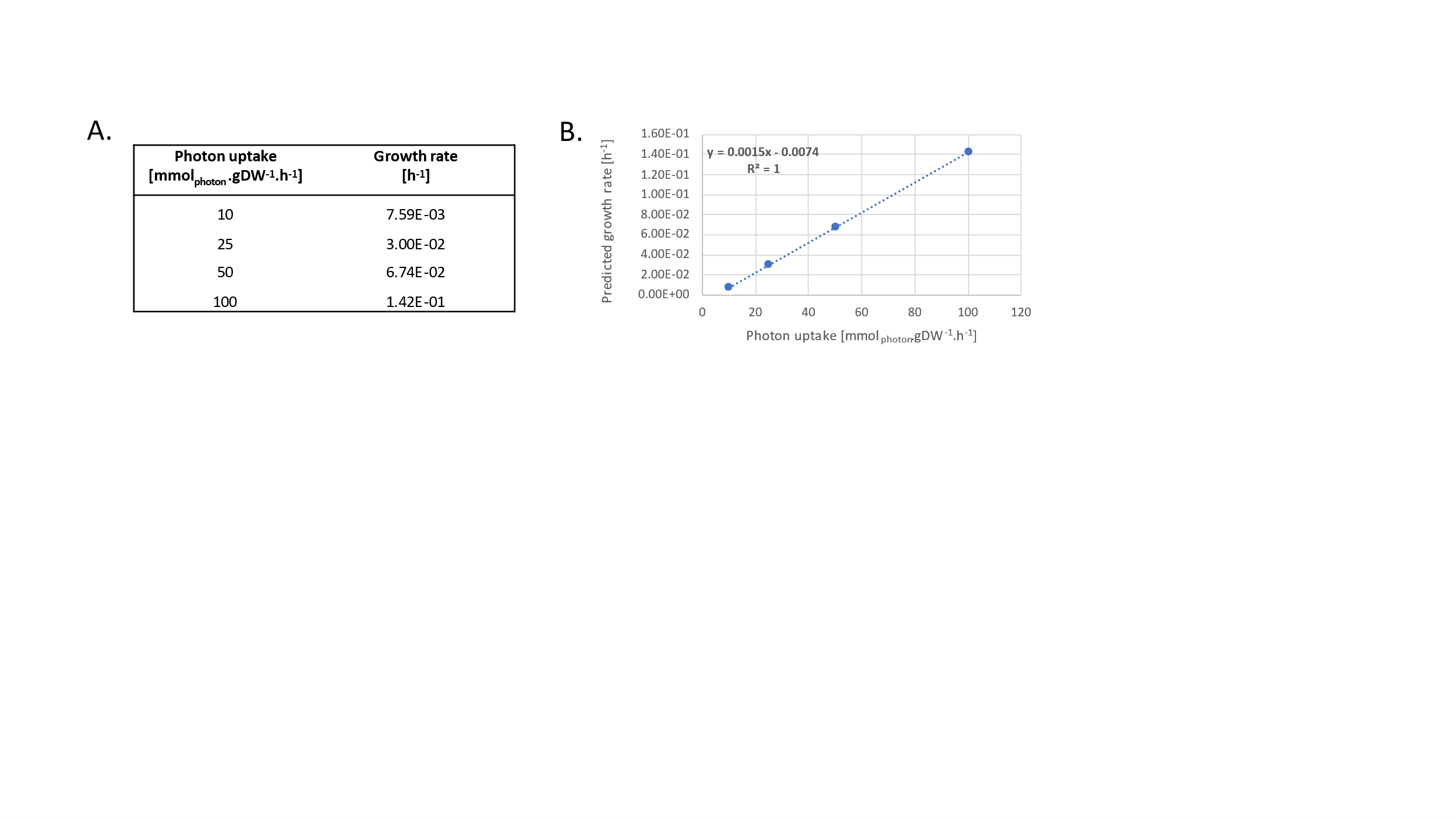

**Figure S5. Linear correlation between photon uptake and optimal growth rate predicted by pFBA.**

Based on this linear correlation, prediction of a growth rate of 0.021 h^-1^ requires a photon uptake of 19 mmol_photon_.gDW^-1^.h^-1^. In consequence, iMgadit23 EX_photon_abs lower bound was set to 19, and pFBA with maximization of WT BOF as objective function was conducted with and without PSQUANTUM pseudo-reaction. Quantum yield (QY) in mol_CO2_.mol_photons_abs_^-1^ was calculated as:

$$QY= \frac{EX\_co2_{\_}flux}{EX\_photon\_abs\_ flux}$$

Biomass composition

**Macromolecules stoichiometric coefficients**

Cultivation of *M. gaditana* cells

For DNA and protein analyses, *M. gaditana* *CCMP52* was cultivated in artificial seawater (ESAW) using 10 times enriched nitrogen and phosphate sources (5.49 10^-3^ M NaNO_3_ and 2.24 10^-4^ NaH_2_PO_4,_ (Dolch *et al.*, 2017)). A volume of 80 mL liquid medium was inoculated at 2.5 10^6^ cells.mL^-1^ and cultivation was achieved in small photobioreactors (Multi-Cultivator MC 1000, Photon Systems Instruments, Czech Republic) at 24°C, in continuous light (illumination at 60 μmol_photon_.m^-2^.s^-1^). Culture mixing was achieved by gas bubbling in which CO_2_ levels were maintained constant at 0.5% as in air-lift photobioreactors. Precise and constant level of CO_2_ was supplied by a Gas Mixing System GMS 150 (Photon Systems Instruments, Czech Republic) following manufacturer’s instructions.

For pigment composition*, M. gaditana CCMP526* was cultivated in GB medium containing anhydrous salts (solution I : 10 mM NaHCO_3_, 445 mM NaCl, 3.5 mM Na_2_SO_4_), hydrated salts (solution II : 3 mM MgSO_4_·7H_2_O, 2.5 mM CaCl_2_·6H_2_O), nitrate and phosphate sources (respectively 33.6 mM KNO_3_ and 2.5 mM K_2_HPO_4_), iron (28 µM Fe-EDTA), trace metals (80 µM Na_2_EDTA·2H_2_O, 4 µM ZnSO_4_·7H_2_O, 1.2 µM CoCl_2_·6H_2_O, 19 µM MnCl_2_·4H_2_O, 0.1 µM Na_2_MoO_4_·2H_2_O, 1.3 µM CuSO_4_·5H_2_O) and vitamins (0.1 µM vitamin H, 0.1 µM vitamin B12, 3.3 µM vitamin B1), pH was set by addition of 100 mM HEPES buffer pH 7.5. A volume of 1.8 L liquid medium was inoculated at 2.6 10^7^ cells.mL^-1^ and cultivation was achieved in ﬂat panel photobioreactors (Labfors 5 Lux, Infors HT, Switzerland) at 26°C under a day:night cycle of 16:8 hours. The incident light intensity increased daily, by keeping the outgoing light intensity 30-40 μmol_photon_.m^-2^.s^-1^, up to 636 μmol_photon_.m^-2^.s^-1^. Culture was mixed by ﬁlter sterilized air with 2% CO_2_ with a ﬂow rate of 1 L.min^-1^.

Pigments extraction and dosage

Concentration of photosynthetic pigments in *M. gaditana* grown under the aforementioned conditions were estimated spectrometrically in terms of mass of pigments per unit volume of suspension (µg/mL). A volume of 1 mL of culture was first centrifuged at 13,400 rpm for 10 min. The medium was discarded and the cells were resuspended by vortexing in 1.5 mL pure methanol. Pigments were extracted for 45min at 44°C in the dark and the extract was centrifuged at 13,400 rpm for 10 min. The optical density OD_λ_ of the supernatant, was measured at wavelengths 750, 665, 652, and 480 nm using a spectrophotometer (BioSpectrometer, Eppendorf). Chlorophyll a and PhotoProtective Carotenoïds (PPC) were estimated according to the correlations:

$$C_{chla} = 13.9 \left( {DO}_{665}- {DO}_{750} \right) {. V}_{2} .\mathcal{l}^{-1}{.V}_{1}^{-1} (Eq.1)$$

$$C_{PPC} = \left( 4 \left( {DO}_{480}- {DO}_{750} \right) \right) . V_{2} .\mathcal{l}^{-1}.V_{1}^{-1} (Eq.2)$$

where V_1_ and V_2_ are respectively volumes of culture and solvent (methanol), ℓ is the thickness of the cuvette used to measure the optical density.

Proteins extraction and dosage

Proteins extraction was performed by adding 100 µL of Lysis buffer composed of Tris HCl 50 mM pH 6,8, SDS 4%, protease inhibitor (complete, Roche Diagnostics) to 2.10^8^ *M. gaditana* cells harvested by centrifugation at 3,500 g for 10 minutes. The mix was nitrogen frozen and thawed 3 times before incubation for 30 minutes at room temperature. For protein precipitation 5 volumes (v/v) of 4°C 80% acetone was added to the mix. After 2 hours at -20°C, the proteins were collected by centrifugation at 13,000 g for 15 minutes at 4°C and the pellet rinsed once with cold acetone 80% before to be dried for 30 minutes at RT. Finally, proteins were resuspended in 100 µL of Lysis buffer. For protein dosage a BCA protein assay kit (BioVision) was used following supplier instructions, bovine serum albumin was used for establishing the standard curve.

DNA extraction and dosage

Cell lysis was run by vortexing a 8.10^8^ cells pellet collected by centrifugation (10 minutes at 3,500 g) into 500 µL of lysis buffer (250 mM Tris pH8.2, 100 mM EDTA, 2% SDS, 100mM NaCl). The solution was then incubated at 60°C for 15 minutes before being vortexed again. DNA extraction was then performed by a classical phenol:chloroform:isoamyl alcohol protocol (25:24:1) used at 1:1 (v/v) and washed twice with chloroform. DNA was precipitated with 30 µL Sodium-Acetate 3M pH5 and 1.5 volume of absolute Ethanol (v/v). After centrifugation and drying, the pellet was resuspended in 30 µL of MilliQ water for photospectrometric dosage (Biospectrometer, Eppendorf).

Lipids, carbohydrates and RNA coefficients

Total lipid, carbohydrate and RNA coefficients were obtained from literature. Lipid and carbohydrates coefficients were calculated based on *M. gaditana* data respectively reported by (Billey *et al.*, 2021) and (Billey *et al.*, 2021; Janssen *et al.*, 2018). RNA coefficient was calculated from *Ochrophyta* data reviewed by (Finkel *et al.*, 2016).

**Specific metabolites coefficients**

DNA composition was calculated from %GC of *M. gaditana* genome. RNA composition was calculated from rRNA sequences of *M. gaditana* genome. Detailed carbohydrate composition was based on (Radakovits *et al.*, 2012; Scholz *et al.*, 2014; Shah *et al.*, 2017). Detailed pigment composition was calculated from (Simionato *et al.*, 2013). Amino acid composition of proteins was calculated based on data from (Lamminen *et al.*, 2019; Medina *et al.*, 2015; Tibbetts, Bjornsson and McGinn, 2015; Volkman *et al.*, 1993). Finally, lipidomic profiles in terms of glycerolipid classes and acyl-chains was detailed based on experimental characterization “untransformed WT” and “MgACSBG#31” strains from (Billey *et al.*, 2021).

**ATP maintenance**

ATP requirements for DNA, RNA, proteins and pigments formation were respectively set to 1.372, 0.4, 4.306 and 2.0 mol ATP per mol of macromolecule (Fachet *et al.*, 2020; Kliphuis *et al.*, 2012; Neidhart F. C., 1996; Oliveira, Nielsen and Förster, 2005). The Growth Associated Maintenance (GAM) requirement was based on *Chlamydomonas reinhardtii* GEM (29.89 mmol_ATP_.g_DW_^-1^.h^-1^) (Boyle and Morgan, 2009). Energetic requirement for non-growth associated maintenance (NGAM), was added to iMgadit23 as an independent reaction (NGAM_c) with a lower_bound constrained of 2.2 mmol_ATP_.g_DW_^-1^.h^-1^ based on *Nannochloropsis* sp. data (Shah *et al.*, 2017; Zhang *et al.*, 2014).

Detailed biomass composition, original data and sources and final stochiometric coefficients are detailed in Supplemental Tables S1 and S2.

Flux simulation

**Default medium composition**

Minimum defined medium, representing photoautotrophic growth with NO_3_ as nitrogen source was used. Unlimited uptake rates of CO_2_, O_2_, H_2_O, protons, phosphate, sulfates, magnesium, mannose, and 6-deoxy-galactose were allowed. This entailed setting the lower bounds of the corresponding exchange reactions to − 100000, as we used the usual convention of writing the exchange reactions in such way that production corresponds to positive fluxes and consumption to negative ones. Complete default constraints are listed in Table S7. In default medium conditions photon uptake was limited to 23 mmol_photon_.gDW^-1^.h^-1^ resulting in a maximum growth predicted with pFBA of 0.027 h^-1^ to be consistent with average growth rate reported in literature for *M. gaditana* (Gentile and Blanch, 2001; Kareya *et al.*, 2020; Kim *et al.*, 2014; Ren and Ogden, 2014).

**Table S6. Default constraints within iMgadit23.**

| **Reaction ID** | **Lower bound** | **Upper bound** |
| --- | --- | --- |
| EX_5mta_ | 0 | 100000 |
| EX_ac_ | 0 | 100000 |
| EX_ade_ | 0 | 100000 |
| EX_arg__L_ | 0 | 100000 |
| EX_ca2_ | 0 | 100000 |
| EX_chol_ | 0 | 100000 |
| EX_co2_ | -100000 | 100000 |
| EX_cobalt2_ | 0 | 100000 |
| EX_fe2_ | 0 | 100000 |
| EX_for_ | 0 | 100000 |
| EX_fru_ | 0 | 100000 |
| EX_fuc__L_ | -100000 | 0 |
| EX_fum_ | 0 | 100000 |
| EX_gal_ | 0 | 100000 |
| EX_glc__D_ | 0 | 100000 |
| EX_gln__L_ | 0 | 100000 |
| EX_glu__L_ | 0 | 100000 |
| EX_glyc_ | 0 | 100000 |
| EX_glyc3p_ | 0 | 100000 |
| EX_gua_ | 0 | 100000 |
| EX_h_ | -100000 | 100000 |
| EX_h2o_ | -100000 | 100000 |
| EX_hco3_ | 0 | 100000 |
| EX_his__L_ | 0 | 100000 |
| EX_inost_ | 0 | 100000 |
| EX_leu__L_ | 0 | 100000 |
| EX_lys__L_ | 0 | 100000 |
| EX_man_ | -100000 | 0 |
| EX_mg2_ | -100000 | 0 |
| EX_na1_ | 0 | 100000 |
| EX_nh4_ | 0 | 0 |
| EX_no2_ | 0 | 100000 |
| EX_no3_ | -100000 | 100000 |
| EX_o2_ | -100000 | 100000 |
| EX_phe__L_ | 0 | 100000 |
| EX_photon_abs_ | -23 | 0 |
| EX_pi_ | -100000 | 100000 |
| EX_ppi_ | 0 | 100000 |
| EX_pro__L_ | 0 | 100000 |
| EX_sbt__D_ | 0 | 100000 |
| EX_so3_ | -100000 | 100000 |
| EX_so4_ | -100000 | 100000 |
| EX_trp__L_ | 0 | 100000 |
| EX_tyr__L_ | 0 | 100000 |
| EX_urea_ | 0 | 100000 |
| EX_val__L_ | 0 | 100000 |
| EX_xylt_ | 0 | 100000 |

**Photon uptake lower bound in differential flux analysis:**

To constrain the photon uptake in the differential flux analysis, the experimental maximum number of photons received by the microalgae was estimated as:

$\frac{a_{s}}{C_{x}}.q_{0}$ = 331 mmol_photon_.gDW^-1^. h^-1^

Where $a_{s}$ is the surface/volume ratio of the photobioreactor ($a_{s}$=73.5m^-1^), $C_{x}$ is the inoculum concentration of microalgae ($C_{x}$ = 48 gDW.m^-3^) and $q_{0}$ is the incident light ($q_{0}$= 216 mmol_photon_.m^-2^.h^-1^).

**Obtention of Rafay *et al.*, 2020 data for quantitative validation**

WebPlotDigitizer (<https://apps.automeris.io/wpd/>) was used to extract volatile suspended solids (VSS; gVSS.L^-1^) and fixed suspended solids (FSS; gFSS.L^-1^) from (Rafay *et al.*, 2020; Figure 3). Dry weight (DW; gDW. L^-1^) was calculated as:

$$DW= VSS+FSS$$

Linear correlation was predicted between DW and VSS (r^2^ = 0.99), with:

$$DW= 0.89 VSS$$

This relation was used to convert maximum specific nitrate uptake rate (mg_NO3_.gVSS^-1^.d^-1^) into unit required to constrain iMgadit23 uptake rate: mmol_NO3_.gDW^-1^.h^-1^ (Table S7).

**T****able S7. Nitrate uptake data used to constrain iMgadit23 for quantitative validation.**

| **Initial nitrate concentration** (mg.L^-1^) (Rafay et al., 2020) | **Nitrate uptake rate**  (mg_NO3_.gVSS^-1^.d^-1^) (Rafay et al., 2020) | **Nitrate uptake rate**  (mmol_NO3_.gDW^-1^.h^-1^) |
| --- | --- | --- |
| 100 | 223 | 1.33E-01 |
| 40 | 199 | 1.19E-01 |
| 56 | 208 | 1.24E-01 |
| **avg** | **210** | **1.25E-01** |
| std | 12.12 | 7.24E-03 |

Reference list

Billey, E. *et al.* (2021) ‘Characterization of the Bubblegum acyl-CoA synthetase of Microchloropsis gaditana’, *Plant Physiology*, 185(3), pp. 815–835. doi: 10.1093/plphys/kiaa110

Boyle, N.R. and Morgan, J.A. (2009) ‘Flux balance analysis of primary metabolism in Chlamydomonas reinhardtii’, *BMC Systems Biology*, 3, p. 4. doi: 10.1186/1752-0509-3-4

Dolch, L.-J. *et al.* (2017) ‘Nitric Oxide Mediates Nitrite-Sensing and Acclimation and Triggers a Remodeling of Lipids’, *Plant Physiology*, 175(3), pp. 1407–1423. doi: 10.1104/pp.17.01042

Fachet, M. *et al.* (2020) ‘Reconstruction and analysis of a carbon-core metabolic network for Dunaliella salina’, *BMC bioinformatics*, 21(1), p. 313. doi: 10.1186/s12859-019-3325-0

Finkel, Z.V. *et al.* (2016) ‘Phylogenetic Diversity in the Macromolecular Composition of Microalgae’, *PloS One*, 11(5), e0155977. doi: 10.1371/journal.pone.0155977

Gentile, M.P. and Blanch, H.W. (2001) ‘Physiology and xanthophyll cycle activity of Nannochloropsis gaditana’, *Biotechnology and Bioengineering*, 75(1), pp. 1–12. doi: 10.1002/bit.1158

Hoffmann, D.Y. and Shachar-Hill, Y. (2023) ‘Do betaine lipids replace phosphatidylcholine as fatty acid editing hubs in microalgae?’ *Frontiers in Plant Science*, 14, p. 1077347. doi: 10.3389/fpls.2023.1077347

Janssen, J.H. *et al.* (2018) ‘Effect of nitrogen addition on lipid productivity of nitrogen starved Nannochloropsis gaditana’, *Algal Research*, 33, pp. 125–132. doi: 10.1016/j.algal.2018.05.009

Kareya, M.S. *et al.* (2020) ‘Photosynthetic Carbon Partitioning and Metabolic Regulation in Response to Very-Low and High CO2 in Microchloropsis gaditana NIES 2587’, *Frontiers in Plant Science*, 11, p. 981. doi: 10.3389/fpls.2020.00981

Kim, C.W. *et al.* (2014) ‘Effect of monochromatic illumination on lipid accumulation of Nannochloropsis gaditana under continuous cultivation’, *Bioresource Technology*, 159, pp. 30–35. doi: 10.1016/j.biortech.2014.02.024

Kliphuis, A.M.J. *et al.* (2012) ‘Metabolic modeling of Chlamydomonas reinhardtii: energy requirements for photoautotrophic growth and maintenance’, *Journal of Applied Phycology*, 24(2), pp. 253–266. doi: 10.1007/s10811-011-9674-3

Lamminen, M. *et al.* (2019) ‘Different microalgae species as a substitutive protein feed for soya bean meal in grass silage based dairy cow diets’, *Animal Feed Science and Technology*, 247, pp. 112–126. doi: 10.1016/j.anifeedsci.2018.11.005

Li, X., Benning, C. and Kuo, M.-H. (2012) ‘Rapid triacylglycerol turnover in Chlamydomonas reinhardtii requires a lipase with broad substrate specificity’, *Eukaryotic Cell*, 11(12), pp. 1451–1462. doi: 10.1128/EC.00268-12

Malitsky, S. *et al.* (2016) ‘Viral infection of the marine alga Emiliania huxleyi triggers lipidome remodeling and induces the production of highly saturated triacylglycerol’, *The New Phytologist*, 210(1), pp. 88–96. doi: 10.1111/nph.13852

Medina, C. *et al.* (2015) ‘Protein Fractions with Techno-Functional and Antioxidant Properties from *Nannochloropsis gaditana* Microalgal Biomass’, *Journal of Biobased Materials and Bioenergy*, 9(4), pp. 417–425. doi: 10.1166/jbmb.2015.1534

Neidhart F. C. (1996) ‘Escherichia coli and Salmonella typhimurium.’, *Cellular and Molecular Biology*, 1, p. 1225. Available at: https://ci.nii.ac.jp/naid/10005158520/.

Oliveira, A.P., Nielsen, J. and Förster, J. (2005) ‘Modeling Lactococcus lactis using a genome-scale flux model’, *BMC Microbiology*, 5, p. 39. doi: 10.1186/1471-2180-5-39

Radakovits, R. *et al.* (2012) ‘Draft genome sequence and genetic transformation of the oleaginous alga Nannochloropis gaditana’, *Nature Communications*, 3, p. 686. doi: 10.1038/ncomms1688

Rafay, R. *et al.* (2020) ‘Growth and Nitrate Uptake in Nannochloropsis gaditana and Tetraselmis chuii Cultures Grown in Sequential Batch Reactors’, *Frontiers in Marine Science*, 7 (9pp). doi: 10.3389/fmars.2020.00077

Ren, M. and Ogden, K. (2014) ‘Cultivation of Nannochloropsis gaditana on mixtures of nitrogen sources’, *Environmental Progress & Sustainable Energy*, 33(2), pp. 551–555. doi: 10.1002/ep.11818

Scholz, M.J. *et al.* (2014) ‘Ultrastructure and composition of the Nannochloropsis gaditana cell wall’, *Eukaryotic Cell*, 13(11), pp. 1450–1464. doi: 10.1128/EC.00183-14

Shah, A.R. *et al.* (2017) ‘Reconstruction and analysis of a genome-scale metabolic model of Nannochloropsis gaditana’, *Algal Research*, 26, pp. 354–364. doi: 10.1016/j.algal.2017.08.014

Simionato, D. *et al.* (2013) ‘The response of Nannochloropsis gaditana to nitrogen starvation includes de novo biosynthesis of triacylglycerols, a decrease of chloroplast galactolipids, and reorganization of the photosynthetic apparatus’, *Eukaryotic Cell*, 12(5), pp. 665–676. doi: 10.1128/EC.00363-12

Tibbetts, S.M., Bjornsson, W.J. and McGinn, P.J. (2015) ‘Biochemical composition and amino acid profiles of Nannochloropsis granulata algal biomass before and after supercritical fluid CO2 extraction at two processing temperatures’, *Animal Feed Science and Technology*, 204, pp. 62–71. doi: 10.1016/j.anifeedsci.2015.04.006

Volkman, J. *et al.* (1993) ‘The biochemical composition of marine microalgae from the class eustigmatophyceae’, *Phycologia*, 29, pp. 69–78.

Zhang, D. *et al.* (2014) ‘On-line modeling intracellular carbon and energy metabolism of Nannochloropsis sp. in nitrogen-repletion and nitrogen-limitation cultures’, *Bioresource Technology*, 164, pp. 86–92. doi: 10.1016/j.biortech.2014.04.083
