## Supplemental Results for "A new genome-scale metabolic model of oleaginous microalgae with refined lipid metabolism elucidates *Microchloropsis gaditana* mutant phenotypes"

### Photosynthesis modeling

Distinction between "absorbed" and "LEF" photons was introduced in iMgadit23 by the addition of PSQUANTUM_h pseudo reaction: 10 photon_abs_h <=> 8 photon_lef_h. pFBA simulations were performed to analyze impact of this reaction on model predictions.

PSQUANTUM pseudo reaction was modified with absorbed/LEF values (8/8, 9/8, 10/8, 11/8 and 12/8), resulting in photon/ATP ratio of 8/3, 9/3, 10/3, 11/3 and 12/3, and pFBA approach was applied with WT BOF maximization as objective function for different photon uptake rates ranging from 0 up to 25 mmol_photon_abs_.gDW^-1^.h^-1^. Similar impacts on growth rate, ATP production (ATPSYN_h reaction flux) and NADPH production (FDXO_h reaction flux) were predicted (Figure S1 B-D).

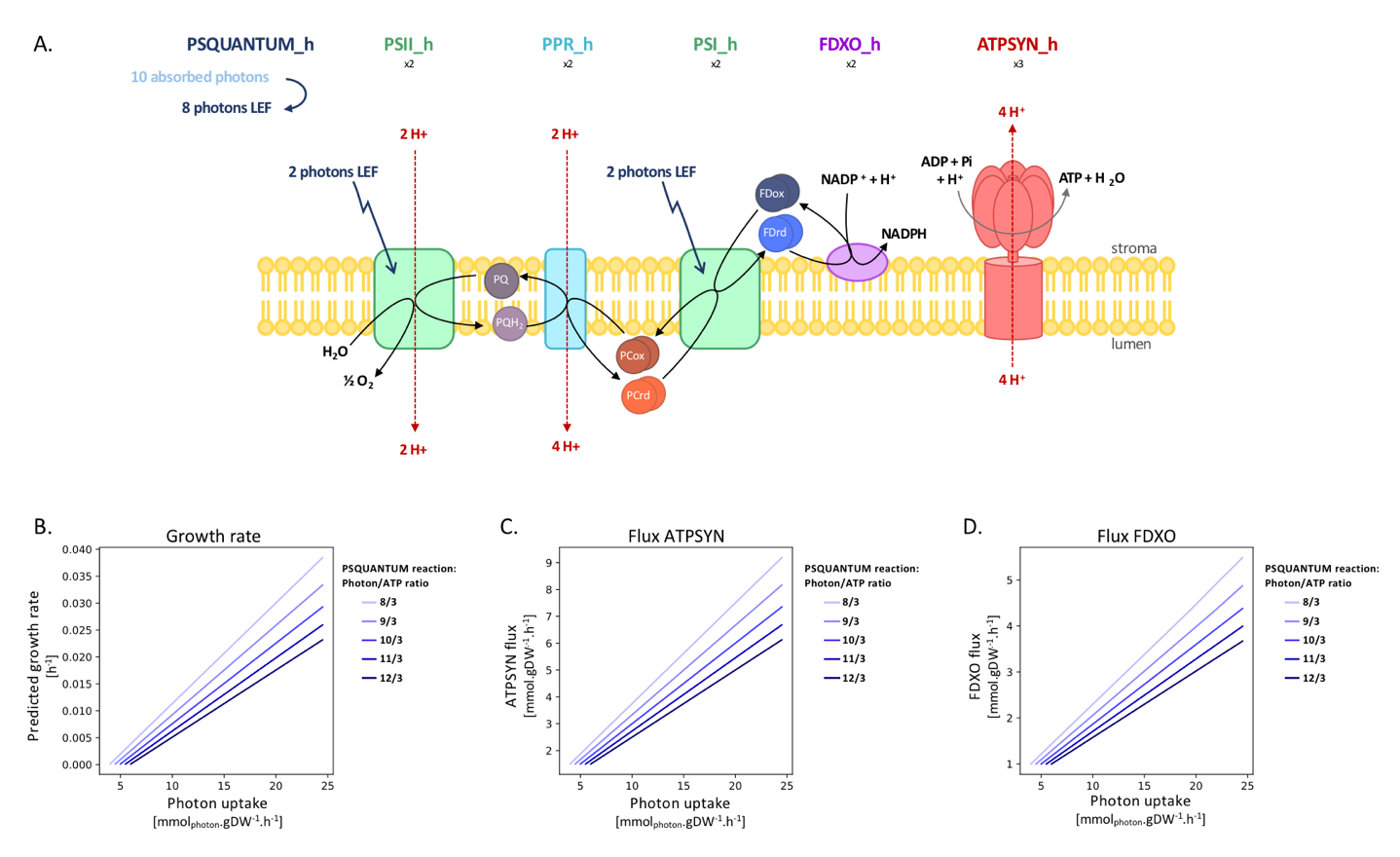

Figure S1. Photosynthesis modeling and impact of PSQUANTUM in iMgadit23.

(A) Light phase of photosynthesis is modeled by six reactions. Photosystems I and II (PSI_h and PSII_h) respectively use two productive photons inducing electron flow through plastoquinones/plastocyanins and ferredoxins (Plastoquinol/Plastocyanin Reductase (PPR_h), Ferredoxin oxidoreductase (FDXO_h)) to produce one molecule of NADPH. The proton gradient established on either side of the thylakoid membrane allows production of ATP by the ATP synthase (ATPSYN_h). PSQUANTUM artificially converts absorbed photons to photons LEF involved in PSI_h and PSII_h reactions, with a ratio of 10 absorbed photons required to produce 8 LEF photons. Protons and electrons flows are represented respectively in red and black. Impact of absorbed photon/ATP ratio due to PSQUANTUM reaction on maximum growth rate (A), ATP production (B) and NADPH production (C) predicted by pFBA.

### Differential flux distribution analysis between WT and *acsbg* strains

#### pFBA

Table S1. log2(pFBA_acsbg/pFBA_WT) results.

| **log2(pFBA_acsbg/pFBA_WT)** | **Number of reactions** | **% of reactions with flux (770)** |
| --- | --- | --- |
| Both off | 1526 |  |
| log2(FC) **< -0.2630** | 606 | 78.70 |
| OFF | 36 | 4.68 |
| log2(FC) in [-0.2630;0.2630] | 38 | 4.94 |
| log2(FC) **>0.2630** | 50 | 6.49 |
| ON | 40 | 5.19 |
| Change sign: positive to negative | 0 | 0 |
| Change sign: negative to positive | 0 | 0 |

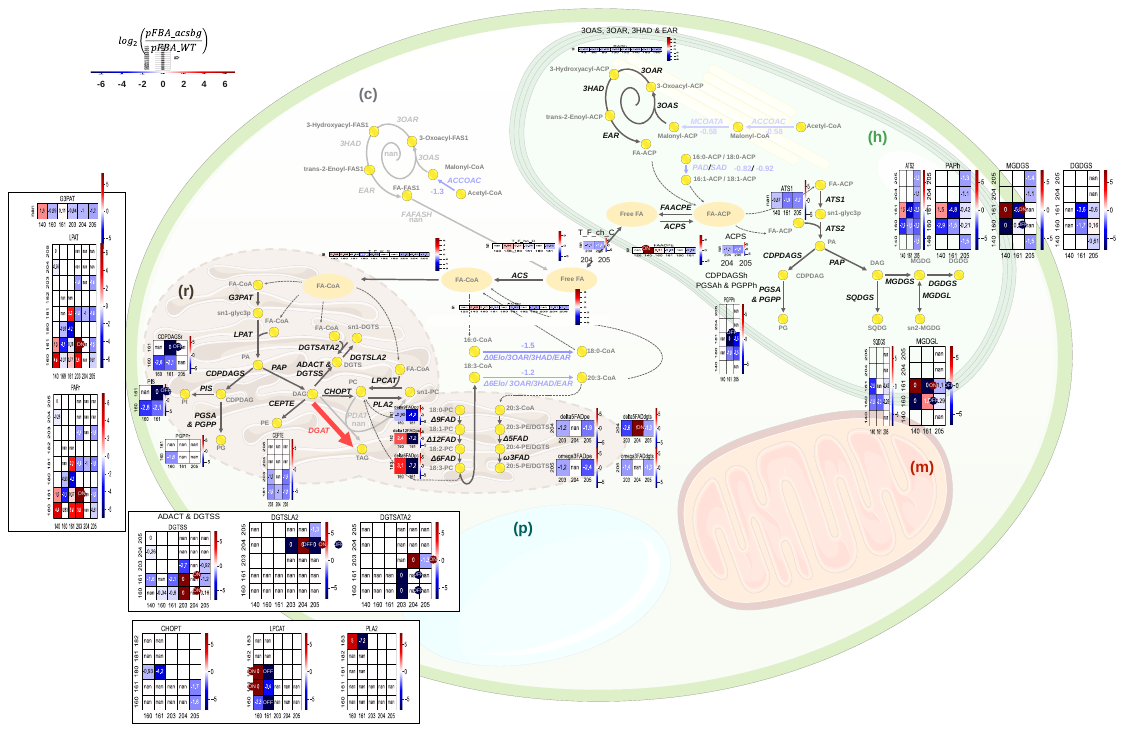

Figure S2. Results of pFBA differential flux analysis for lipid metabolism.

For each reaction, results are shown for all sn-1 (x-axis) and sn-2 (y-axis) couples. For each specific reaction, log2(pFBA_acsbg/pFBA_WT) value is indicated. Red colors indicate reactions upregulated in acsbg. Blue colors indicate reactions downregulated in acsbg. "nan" indicates that reaction was predicted to have no flux both in pFBA_acsbg and pFBA_WT. Finally, empty value indicates that reaction does not exist in iMgadit23.

#### MOMA

Table S2. log2(MOMA_acsbg/pFBA_WT) results.

| **log2(MOMA_acsbg/pFBA_WT)** | **Number of reactions** | **% of reactions with flux (1009)** |
| --- | --- | --- |
| Both off | 1287 |  |
| log2(FC) **< -0.2630** | 373 | 36.97 |
| OFF | 16 | 1.59 |
| log2(FC) in [-0.2630;0.2630] | 253 | 25.07 |
| log2(FC) **>0.2630** | 80 | 7.93 |
| ON | 279 | 27.65 |
| Change sign: positive to negative | 5 | 0.50 |
| Change sign: negative to positive | 3 | 0.30 |

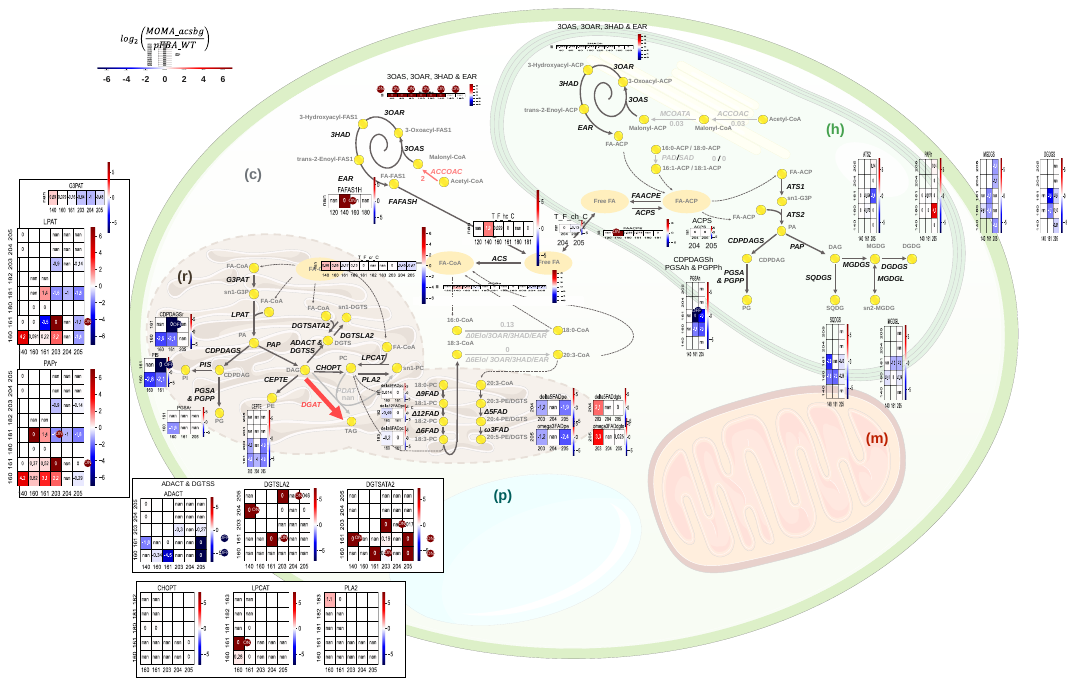

Figure S3. Results of MOMA differential flux analysis for lipid metabolism.

For each reaction, results are shown for all sn-1 (x-axis) and sn-2 (y-axis) couples. For each specific reaction, log2(MOMA_acsbg/pFBA_WT) value is indicated. Red colors indicate reactions upregulated in acsbg. Blue colors indicate reactions downregulated in acsbg. "nan" indicates that reaction was predicted to have no flux both in MOMA_acsbg and pFBA_WT. Finally, empty value indicates that reaction does not exist in iMgadit23.
